# Conflicting roles of cell geometry, microtubule deflection and orientation-dependent dynamic instability in cortical array organization

**DOI:** 10.1101/2024.09.07.611822

**Authors:** Tim Y.Y. Tian, Geoffrey O. Wasteneys, Colin B. Macdonald, Eric N. Cytrynbaum

## Abstract

The self-organization of cortical microtubule arrays within plant cells is an emergent phenomenon with important consequences for the synthesis of the cell wall, cell shape, and subsequently the structure of plants. Mathematical modelling and experiments have elucidated the underlying processes involved. There has been recent interest in the influence of geometric cues on array orientation, be it direct (cell shape) or indirect (tension in the membrane). However, the mechanical influence of membrane curvature on these elastic filaments has largely been ignored. A previous model was proposed to describe how the anchoring process may control the deflection of individual microtubules seeking to minimize bending on a cylindrical cell. We incorporate this process into a model of interacting microtubules and find the cell curvature influence to be significant: the array favours orientations parallel to the direction of elongation rather than the expected transverse direction. Even without elasticity, the geometry of large cells hinders robust microtubule organization. These results suggest the necessity of additional processes to overcome these factors. We propose an orientation-dependent catastrophe rate, hypothetically caused by cellulose microfibrils impeding microtubule polymerization. We find a combination of anchoring and impedance to be sufficient to generate transverse arrays despite the geometric influences.

**Significance Statement:** The organization of microtubule (MT) polymers into parallel arrays along the two-dimensional cortex of plant cells is crucial for directional cell growth and plant development. Despite decades of experimentation and more recent computational modelling, understanding the mechanisms that orient cortical MTs remains incomplete. With computational modelling, we have re-examined an assumption common to many models: that MTs grow along straight (geodesic) paths rather than minimizing bending. We model MT bending, and find a significant disruption of transverse MT ordering, especially in larger cells. We find that angle-dependent MT behaviour can counteract the effect of bending in certain contexts.

## Introduction

Plant morphology depends on microscopic-level processes within each cell. In particular, microtubule (MT) polymers organize along the cortex of cells in close association with cellulose deposition [6]. In turn, the orientation and structure of the cellulose microfibrils provides a mechanical constraint determining the direction in which the cells can elongate. Collectively, this contributes to the macroscopic shape of plant organs. The organization of cortical microtubules (CMTs) is of particular interest because plants lack the MT organizing centres present in animal cells. Instead, self-organization is observed to arise from various factors including MT-MT interactions along the cortex [14] (illustrated in panel B of Fig. 1), modes of MT nucleation [9, 36, 26], cell geometry [3, 8, 11], and the regulation of MT-associated proteins involved with these processes [4, 27, 61, 33, 37]. Due to the difficulty of studying each factor *in vivo*, quantitative models have been effective at testing hypotheses relating to each factor in isolation. The most direct method of modelling is particle-based, where the “particles” are MTs represented as curves interacting on the two-dimensional cortex. These models have proven helpful in analyzing CMT organization, and remain a popular tool for exploring the importance of each process [41, 18]. Investigating the fundamental principles behind array organization, these models have shown the importance of dynamic instability parameters (polymerization speed, depolymerization speed, catastrophe rate, etc.), and the various resolutions of MT-MT interactions (entrainment, crossover, collision-induced catastrophe) [47, 2, 53]. Subsequently, more subtle mechanisms such as MT-dependent nucleation (whereby MTs nucleate from existing MTs) and severing (whereby an existing MT is severed) have been implemented to demonstrate their roles in increasing (or decreasing) organization [2, 13, 17, 12]. More recently, models of protoxylem patterning have revisited the role of MT-dependent nucleation where it facilitates banded MT formation[45, 26, 25, 42].

**Figure 1:**
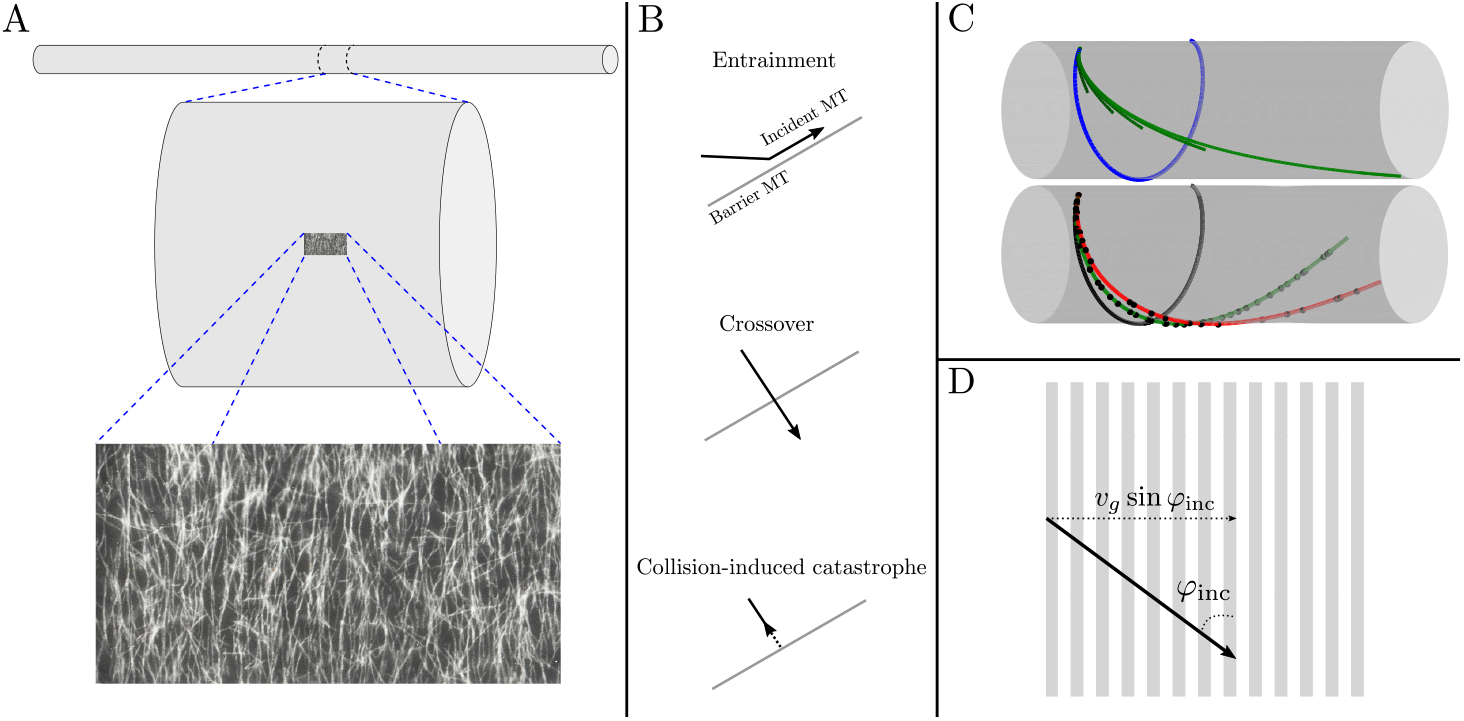
Panel A: An example of cortical microtubule organization transverse to the axis of elongation in an elongating *Nitella tasmanica* internodal cell. From top to bottom: increasing levels of magnification, with features at each level to relative scale. The microscopy image (55*µ*m*×*110*µ*m) was taken at the middle (exact location not measured) of a cell 30.9mm in length and 0.59mm in diameter. Only 12mm of the full length is illustrated at the top level. The image is from an experiment observing repolymerization after oryzalin treatment; methods are described in [60]. Panel B: MT-MT interactions. When a growing incident MT collides with a barrier MT, the incident MT may entrain (also known as zippering or bundling), cross over, or switch to depolymerization (catastrophe). Panel C: The modelling of microtubule deflection on a cylinder. All curves represent MTs with the same initial position and initial growth direction. The MTs are clamped at their base, fixing the initial position and tangential angle and prohibiting lateral movement of the entire MT. Top (C): a comparison of MT shapes that minimize the bending energy Eq. 1 (green) and corresponding geodesic (blue) for MTs anchored and clamped only at one end. Geodesics on a cylinder are helices of a constant pitch, determined by the initial angle of nucleation. In this case, the initial angle is nearly transverse to the long axis. Different shades of green represent the shapes corresponding to energy minimizers for different MT lengths, with the longest green MT having the same length as the blue geodesic MT. Longer MTs tend to deviate more from the geodesic due to the preference to align parallel in the direction of lowest curvature, the longitudinal direction. Bottom (C): Energy-minimizing shapes of three MTs of the same length, using the anchoring model in [51]. Black dots indicate anchoring positions generated with the stochastic model. The geodesic (black) is the infinite rate limit, covering the entirety of the MT in anchors. The green MT is simulated with parameters (cylinder radius, anchoring rate, and polymerization speed) estimated from data (see Table S1), and red with an arbitrarily chosen lower anchoring rate. Panel D: Illustration of a potential model for cellulose feedback. While a MT (solid black) grows at speed *v*_*g*_ with angle of incidence *φ*_inc_, its tip encounters ridges of transversely oriented cellulose microfibrils (grey lines). The rate at which it encounters the ridges is proportional to its horizontal speed *v*_*g*_ sin *φ*_inc_ (dotted arrow).

One important question is how the CMT array in diffusely expanding cells, which expand uniformly along one axis, orders in a manner transverse to the axis of cell elongation during its rapid growth phase. An example of this in giant algae (*Nitella tasmanica*) cells is shown in Fig. 1 A. Previous work [3] found that

MT-stabilizing proteins known as CLASP localizes to the sharp top/bottom edges of root cells in the division zone in *Arabidopsis*, facilitating the passage of MTs around these edges. In CLASP knockouts, MTs in those same cells underwent a transition to depolymerization upon reaching these edges, a process known as edge-induced catastrophe (EIC). These knockouts are observed to have highly ordered transverse arrays not seen in the wild-type case. *In silico* testing of EIC found that it was indeed sufficient to induce transverse arrays [3]. Thus, there was compelling evidence that EIC is responsible for orienting CMT arrays in the transverse direction [18].

To understand the basic factors governing CMT organization, the early models made simplifying assumptions on the cellular geometry and MT trajectories; cells were planar, cylindrical, or polyhedral and MTs grew along straight lines or geodesics, the generalization of straight lines to curved surfaces. We will refer to the curves traced by the MT body as the MT shape. In practice, this results in MTs growing along straight lines [2, 4, 7], or helices along a cylindrical surface [17, 52, 45]. More recent work has incorporated fluctuating MT shapes with finite persistence lengths [35, 25].

Building on these foundations, there has been recent interest in modelling the role of complex cell geometries on MT organization and CMT response to mechanical stress in the cell wall [41, 7, 35, 11, 21, 30]. This poses new modelling challenges: from generalizing previous methods to complicated shapes, to reconsidering the validity of previous model assumptions when applied to this new context. Cell geometry can impact organization in different ways: modifying the connectivity of regions, introducing sharp corners, and the influence of local curvature on MT bending. In the direction of the first two geometric influences, researchers have studied MT growth on complex geometries while keeping with the geodesic assumption [35, 7]. Thus far, all models of interacting MTs ignore the effects of membrane curvature on the bending of these relatively stiff polymers. We explore the question: could curvature effects impact CMT organization? If so, how can this be regulated by the cell?

The effect of curvature was first considered by Lagomarsino et al. [28] in the hypothetical case of individual MTs with lengths on the scale of a whole cylindrical cell. Their predictions of MTs favouring low curvature configurations were generally consistent with their *in vitro* experiments of MTs confined to the volume of microfabricated chambers. *In vivo*, MTs are constrained to the membrane by anchoring proteins, although the exact proteins responsible are largely unknown [21]. In a cylindrical geometry, theory predicts that MTs should favour the longitudinal direction, corresponding to the flattest direction. This is indeed observed in the non-expanding “shank” regions of tip-growing cylindrical cells such as root hairs and pollen tubes in contrast to the transverse CMT arrays typically observed in cells that are elongating by diffuse expansion [58]. A purely bending-energy-minimizing MT would orient and grow parallel to the long axis. If anchored in a direction offset from the long axis, lateral rotation of the entire MT is ruled out. Instead, the laterally unconstrained growing tip would deflect away from the geodesic path prescribed by the initial anchored angle toward the curvature-minimizing direction. The question of how this MT-length dependent deflection compares to the geodesic case was further studied by Bachmann et al. [5]. In particular, it was found that the unanchored MT tip may deviate – bifurcate in special cases – from geodesics if the inter-anchoring distance is large enough. Conversely, the geodesic shape may be preserved in the limit of continuous anchoring. This is seen in the top of Fig. 1 C, where energy minimizers of various lengths are compared to a geodesic curve with the same nucleation angle. Subsequently, a minimal model of anchoring was proposed by Tian et al. [51] to describe MT shapes, and how this may be regulated by mechanics induced by randomly placed anchors. In the model of [51], there are two main factors: the anchoring rate and the total MT length. This is illustrated in the bottom of Fig. 1 C: the larger the anchoring rate (red, green, and black in increasing order), the smaller the MT segments and therefore the smaller the deflections from geodesics. The cumulative effects of many segments may lead to a larger deflection regardless of the small anchoring lengths, making the net effect subtle to predict. This is shown by the final positions of each MT tip. *In vivo*, MTs may vary in length due to dynamic instability and their various interactions. Thus, it remained difficult to deduce whether the bending effect on individual MTs influenced array organization without a full simulation of interacting MTs. Here, we present a simulation of interacting and bending MTs to examine the influence of curvature-induced deflection. We then propose a hypothesis, based on cellulose feedback, for a transverse biasing mechanism that may counteract this bending influence.

## Methods

We first summarize aspects of the model that have been described in previous works, namely, parameters used, and the anchoring process. Lastly, we propose a new model of orientation-dependent catastrophe, in the context of cellulose feedback. Further details of the simulation method and CMT order quantification are found in the Supplementary Information.

### System Parameters

We base our system parameters on those used in previous studies [52, 13, 53] to provide a good basis for comparison. These are idealized or estimated from experimental sources. The dimensional values are presented in Table S1. The dynamic instability parameters are those measured from tobacco BY-2 cells [56], which have a geometry close to that of a cylinder and are diffusely expanding. We used the two-state model of dynamic instability: growth and shrinkage of the plus end with pause states excluded. There is constant depolymerization of the minus end, with the rate given by [46]. For simplicity, we included isotropic nucleation without MT-dependent nucleation. We set the default nucleation rate as in [17]. Severing is absent. The anchoring rate parameter (described later), 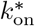, was estimated by Allard et al. [1] in *Arabidopsis* cells.

The domain is initially set such that the cylinder height equals circumference (*L* = 2*πR*). We refer to this as the long cylinder case, because the height is *π* times the diameter. Unravelled, the cylinder surface is square. This allows for a direct study of curvature influence while maintaining a biologically relevant geometry. In addition, we simulate on a shorter cell where the height matches the diameter. Because elastic MTs will tend toward the longitudinal direction, we expect a competition between the longitudinal bias from the bending influence and transverse bias due to EIC. To examine whether EIC remains sufficient in generating transverse CMTs, we implement a strong catastrophe condition: 100% catastrophe along the edges of the cylindrical caps. For this reason, we omit the study of MTs on the cap surfaces.

### Elastic MT Model

As with the minimal anchoring model by Tian et al. [51], we make three fundamental assumptions on the processes relating to anchoring and MT deflection. First, anchoring proteins are abundant and, on the micrometer scale, have no preferred attachment site along MTs or the membrane. Observations of cortical MTs thus far have not shown evidence otherwise. Although there are potential additional predetermined locations of anchor points along the membrane, these are believed to be sparse [21]. In other words, the majority of anchoring locations are likely not predetermined. Second, MTs seek to minimize their bending energy. It is generally accepted that MTs are relatively stiff filaments with large persistence lengths [24], with *in vitro* measurements in the millimeter range from tethering assays [44]. Purely from energy considerations, bending energy generally dominates thermal fluctuations [4]. While factors such as cytoplasmic streaming remain possible additional influences on MT shape [19, 40, 4], these are not considered in this model – as is the case for all models thus far. Third, anchoring proteins clamp the MT in place without imposing additional torque to change the MT orientation from its pre-anchored orientation. The possibility of anchoring proteins exerting additional torque or otherwise altering MT shape remains open to exploration, and may be a hypothetical pathway for MT sensing of membrane tension and/or mechanical stress [21]. This is not considered further in this work.

We model the cell as a finite cylinder with MTs as elastic rods growing constrained to the cell’s cortex, the cylinder surface. In the simplest case, the CMT is fixed at its originating (minus) end, with the growing (plus) end free. The shape of the MT is determined by minimizing the curvature-energy functional,

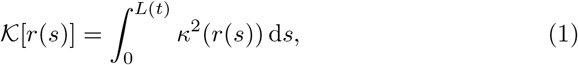

where *r*(*s*) is the arc-length parametrization of the MT body in ℝ^3^, *κ* the curvature, and *L*(*t*) the length of the MT at a given time. The study of such solutions can be found in [5]. Notably, the MT shape depends on the length *L* of the MT at any moment in time. At a time *t*, the solution to the variational problem can be found by expressing *r*(*s*) in terms of the angle of the tangent to MT body relative to the circumferential direction and solving the Euler-Lagrange equations. We denote this parametrization as *φ*(*s*; *t*), where *s* is the arc-length and *t* is the time which encodes the dependence on MT length (*t* = *L/v*_*g*_). This explains how the shapes of inter-anchor segments are determined. Next, we summarize how anchoring is handled in the model [51].

### Anchoring Process

Following [51], anchoring is treated as a stochastic process where anchoring proteins attach randomly along the unanchored growing MT tip, with a rate proportional to its length. The constant of proportionality is denoted as *k*_on_. This results in a non-dimensional parameter,

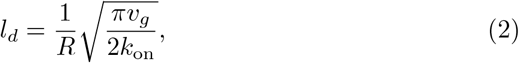

where *R* is the cylinder radius. This is the mean MT tip length prior to the first anchor attachment, the *mean first anchor length*, relative to the radius. This is not the mean length to the first anchor because the anchor can attach anywhere along the free length. As an illustration, we refer to the longest green curve in the top of Fig 1C. Suppose a MT is this length at the instance an anchor protein attaches anywhere along the MT body. *l*_*d*_ corresponds to the (mean) distance from the nucleation point to the tip, rather than the distance from the former point to the new anchor point. For large *l*_*d*_, the MT tip grows longer and has more opportunity to deflect toward the longitudinal curvature-minimizing direction. *l*_*d*_ therefore controls the influence of membrane curvature on individual MTs. The default value 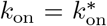 used here corresponds to the rate constant measured by fitting a model to data [1]. Although the aforementioned model is slightly different from ours in that it includes a detachment rate, that rate was found to be relatively small so our implementation is a reasonable estimate. Combined with the other default parameters used, we arrive at *l*_*d*_ = 0.23*R* (4.7*µ*m in dimensional form). This is significantly smaller than the smallest of persistence lengths (the length scale at which thermal fluctuations become significant) measured *in vitro* [44], suggesting that thermal fluctuations should have little impact on the MT shapes prior to anchoring.

We incorporate the elastic MT model into a simulation of interacting MTs with collision resolutions following that of existing models [2, 3, 13, 17, 52, 35, 25, 45, 42], differing in some details. These differences and further details of the simulation method used to incorporate MT bending with MT interactions can be found in the Supplementary Information.

### Orientation-Dependent Catastrophe: Cellulose Feedback

Local influences on CMTs away from the cell edges that are orientation dependent have been suggested (e.g., tension [21, 30]) but mechanistic explanations are still lacking. To investigate a hypothetical mechanism to compete with the longitudinal bias caused by MT deflection, we propose an orientation-dependent catastrophe rate that increases catastrophe along the longitudinal direction. This can be modelled as,

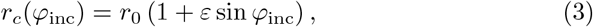

where the catastrophe rate, *r*_*c*_, has an intrinsic rate *r*_0_, and is dependent on the angle of incidence *φ*_inc_ between the MT tip and the transverse axis. The relative strength of the angle dependence is given by *ε*. We note that the functional form of this angle dependence in Eq. 3 may generically describe a phenomenological model of orientation-dependent catastrophe. However, we also provide a hypothetical mechanistic description that can be tested with future experiments.

In particular, this angle dependence could be due to MTs sensing ridges of transversely oriented cellulose microfibrils. This is illustrated in Fig. 1 D. The persistence of transversely oriented microfibrils – even in the absence of well-ordered MTs – is supported by several studies involving high-resolution field-emission scanning electron microscopy [49, 48, 23, 20]. Additionally, experimental work has observed changes in CMT orientation after perturbing the cellulose microfibril ordering by treatment with a cellulose synthesis inhibitor [23]. There was a statistically significant increase in deviation of CMT transverse orientation after the microfibrils lost order [23]. Subsequent studies have shown that mutations affecting cellulose production and quality can alter CMT orientation [10, 29, 34]. This motivates the following self-reinforcing model where cellulose microfibril orientation feeds back into MT orientation.

In this work, we follow the commonly used model of MT dynamic instability in which the switch between states is a Poisson process. In particular, the probability of a polymerizing MT undergoing catastrophe in a small time span Δ*t* is given by *r*_*c*_Δ*t*. Given the high turgor pressure of plant cells, protoplast tension could enable CMTs to sense the cell wall textural landscape [21]. We hypothesize that CMTs encounter ridges along inner face of the membrane, corresponding to cellulose microfibrils, which hinder their polymerization. This may occur by MTs being directly obstructed by the ridges, or indirectly by e.g., protein structures associated with the microfibrils. We assume that a MT growing with an angle of incidence *φ*_inc_ with respect to the cellulose microfibrils has an additional probability of catastrophe that is proportional, with constant of proportionality *λ*, to the number of ridges *m*(Δ*t*) encountered in a time interval Δ*t*. The number of ridges encountered is the product of the ridge density *ρ*, measuring the number of transverse cellulose microfibrils per unit vertical distance and the vertical distance travelled Δ*z* = *v*_*g*_ sin *φ*_inc_Δ*t*. In all, the probability of a MT polymerizing at speed *v*_*g*_ undergoing catastrophe is given by the sum of its intrinsic instability (*r*_0_Δ*t*) and the cellulose feedback factor,

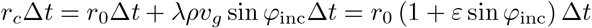

where 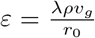. We note that *φ*_inc_ in this case refers to the angle of incidence at the MT tip.

Previous works used a single catastrophe rate, the “observed” catastrophe rate *r*_obs_, under the assumption of isotropic angle-independent dynamic instability. There remains a question of how the previously measured catastrophe rate *r*_obs_ is related to *r*_0_ in Eq. 3. Following the assumption that *r*_obs_ was calculated in the context of a uniform population density of measured MTs with respect to tip orientation (described in Sec. S1.6), our model gives the orientation-dependent catastrophe rate as,

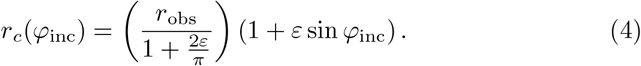

This model may arise as a coarse-grained description of a model that considers the detailed polymerization process. In particular, the constant of proportionality describing the sensitivity to cellulose feedback, *λ* (and consequently *ε*), may depend on physical quantities such as the turgor pressure, diameter of cellulose microfibrils in relation to MT diameter, membrane elasticity, presence of MT-stabilizing proteins or additional structures, etc. It remains to be seen what values are physically realistic.

## Results

To understand the interplay between the transverse-biasing influence of EIC, the longitudinal-biasing influence of curvature-induced MT deflection, and the transverse-biasing influence of orientation-dependent catastrophe, we ran three stages of simulations. The first stage included EIC and curvature-induced MT deflection using the default parameters in Table S1 in the context of a short cell. In the second stage, we investigated the impact of cell geometry by repeating this on a longer cell where we hypothesized that curvature-induced deflection would have a stronger influence. Simultaneously, the anchoring rate *k*_on_ and hence the degree of MT deflection was varied. Finally, we tested whether an orientation-dependent catastrophe rate can robustly generate transverse arrays despite the presence of curvature-induced MT deflection.

Because of the competition between EIC and MT deflection in different parts of the domain, we partitioned the domain into circumferential bands and quantified the angular distributions of CMTs within each band. To quantify MT ordering, we unwrapped the cylinder surface into a rectangle, ignoring the end caps, and computed the nematic order parameters *S*_2_ and |Ω| from the angular distributions of MTs. *S*_2_ measures the degree of ordering where a value of 1 corresponds to perfectly parallel MTs and 0 corresponds completely disordered MTs. Ω measures the dominant angle with respect to the transverse axis with values between *−*90^*°*^ and 90^*°*^. We took the absolute value of this angle. This is further described in Sec. S1.5.

### In Short Cells, EIC Produces Relatively Transverse Arrays

Taking the geodesic limit (*k*_on_ *→ ∞*) in a short cell with EIC, we found strongly transverse arrays (low dominant angle |Ω| and high order *S*_2_) throughout the whole domain, seen in Fig. 2. This is consistent with previous modelling studies that focused on domains with a similar aspect ratio and/or prism-shaped cells with flat faces where MTs travel along straight lines [3, 31]. *In vivo*, this aspect ratio would correspond to that of recently divided cells entering the elongation phase.

**Figure 2:**
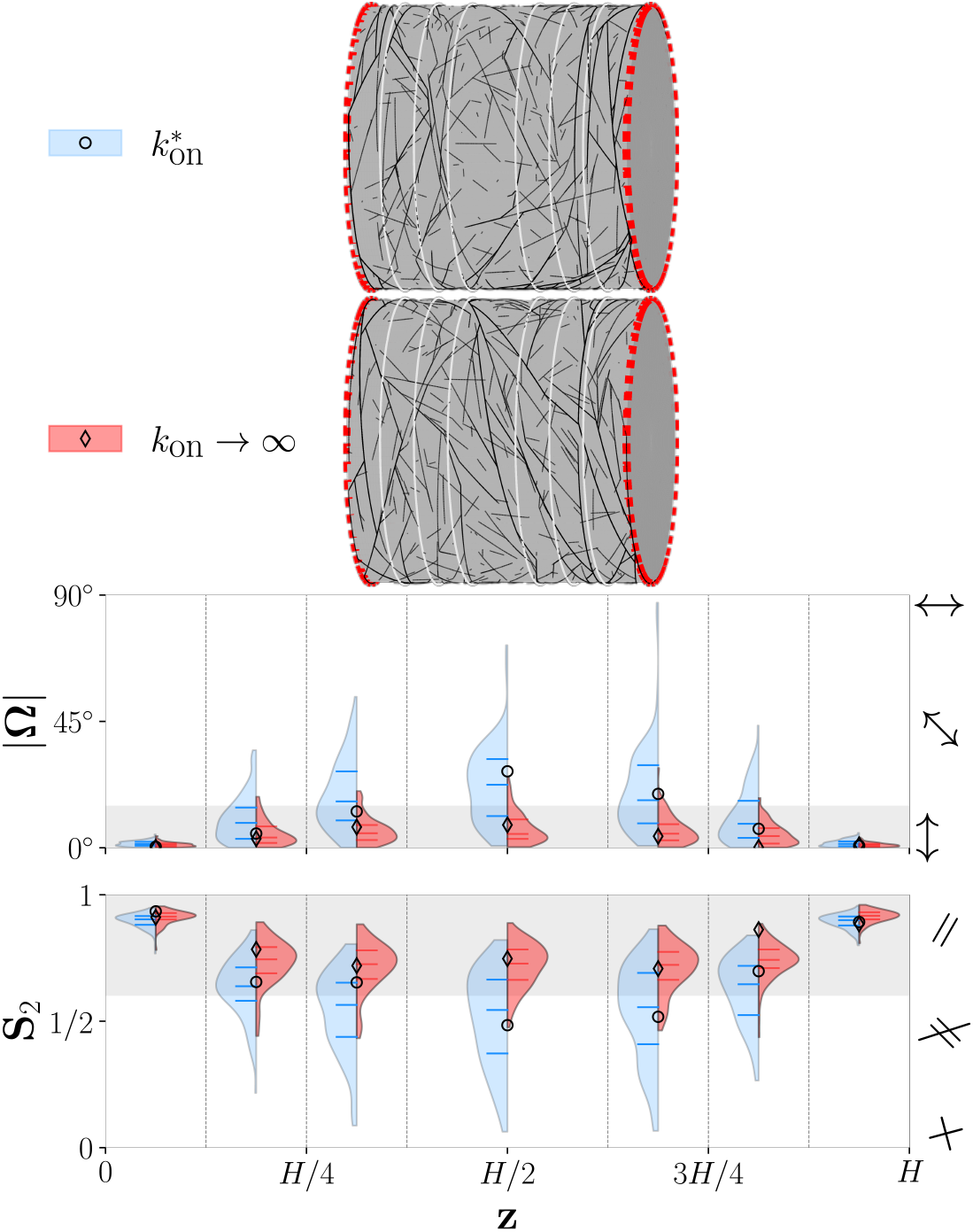
Ordering of the cortical microtubule array in a short cell plotted for two values of the anchoring rate *k*_on_, with all other parameters set as default in Table S1. Top: visualization of individual simulations in short cylindrical cells. Microtubules on the far side of the cylinder are not shown. Red dotted lines indicate catastrophe-inducing edges and white lines indicate the domain partitioning. Anchoring rates used were the experimentally estimated value 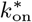, or taking the infinite (geodesic) limit. The corresponding values are labeled on the left of each cylinder. Bottom: violin plots of the order parameters |Ω| (dominant angle) and *S*_2_ (order) plotted along the respective bands of the cylindrical domain set with the long axis horizontal (*z*). The first, second, and third quartiles are indicated in each violin half-plot. Gray-shaded regions provide a qualitative estimate of expected order parameters for transverse microtubule arrays: *S*_2_ *≥* 0.6 [50] and |Ω| *≤* 15^*°*^. Order parameter statistics were calculated by considering the angular distribution for microtubules within each band of the domain after 10h simulation time, and then averaging over 100 simulations. The circle and diamond indicators represent values of the order parameters for the particular simulation snapshots with 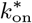 and *k*_on_ *→ ∞* illustrated above.

Next, we introduced curvature-induced deflection by setting the anchoring rate to the experimentally estimated value 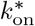. We found the ordering to remain strongly transverse near the edges. However, the MTs in the middle were less ordered and slightly oblique. This raised the question of how to define the CMT as being “sufficiently transverse”. Although there have been measurements of angular distributions in various experiments (e.g., [3, 23, 31]), there does not appear to be sufficient data for a direct quantitative comparison of order parameters as measured *in vivo* in a statistically meaningful sense.

As shown above, transverse MTs appeared in boundary layers localized to the edges. We investigated whether a globally transverse array can form in the presence of MT deflection, even in the case where the boundaries are further apart in the context of a long cell.

### Microtubule Deflection Prevents Global Edge-Induced Transverse Arrays in Long Cells but Anchoring Modulates its Role

To study the role of MT deflection in the case of a long cell and how it can be controlled by anchoring, we varied the anchoring rate *k*_on_ by using multiples of the experimentally estimated value 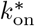 or taking the geodesic limit. This is shown in Fig. 3. We found the longitudinal influence to be strong for the experimentally measured 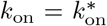. Although there was strong transverse order (low |Ω| and high *S*_2_) of CMTs close to the edges due to EIC, longitudinal deflection dominated (high |Ω| and moderate *S*_2_) toward the middle of the cell. In the transition from the edge toward the middle, this competition between the two influences was seen by the lower *S*_2_ values in the second and third bands from the edges. This observation of transverse ordering only localized to the edges and strong longitudinal ordering elsewhere also occurred for cell lengths intermediate to the cells shown in Figs. 2 & 3 (see Fig. S7). By increasing *k*_on_, thereby decreasing the deflection, the longitudinal influence was mitigated. This is consistent with our previous hypothesis [51] of anchoring being a potential mechanism for controlling array orientation. Fitting the dominant angle curve to a Hill function (Fig. S6), we found that increasing anchoring allows the EIC signal propagate further into the cell in addition to lowering the overall array angle. Whereas the experimentally estimated rate resulted in strong longitudinal ordering, roughly an order of magnitude increase gave ordering similar to the geodesic case. However, the ordering of the geodesic case deserved particular attention.

**Figure 3:**
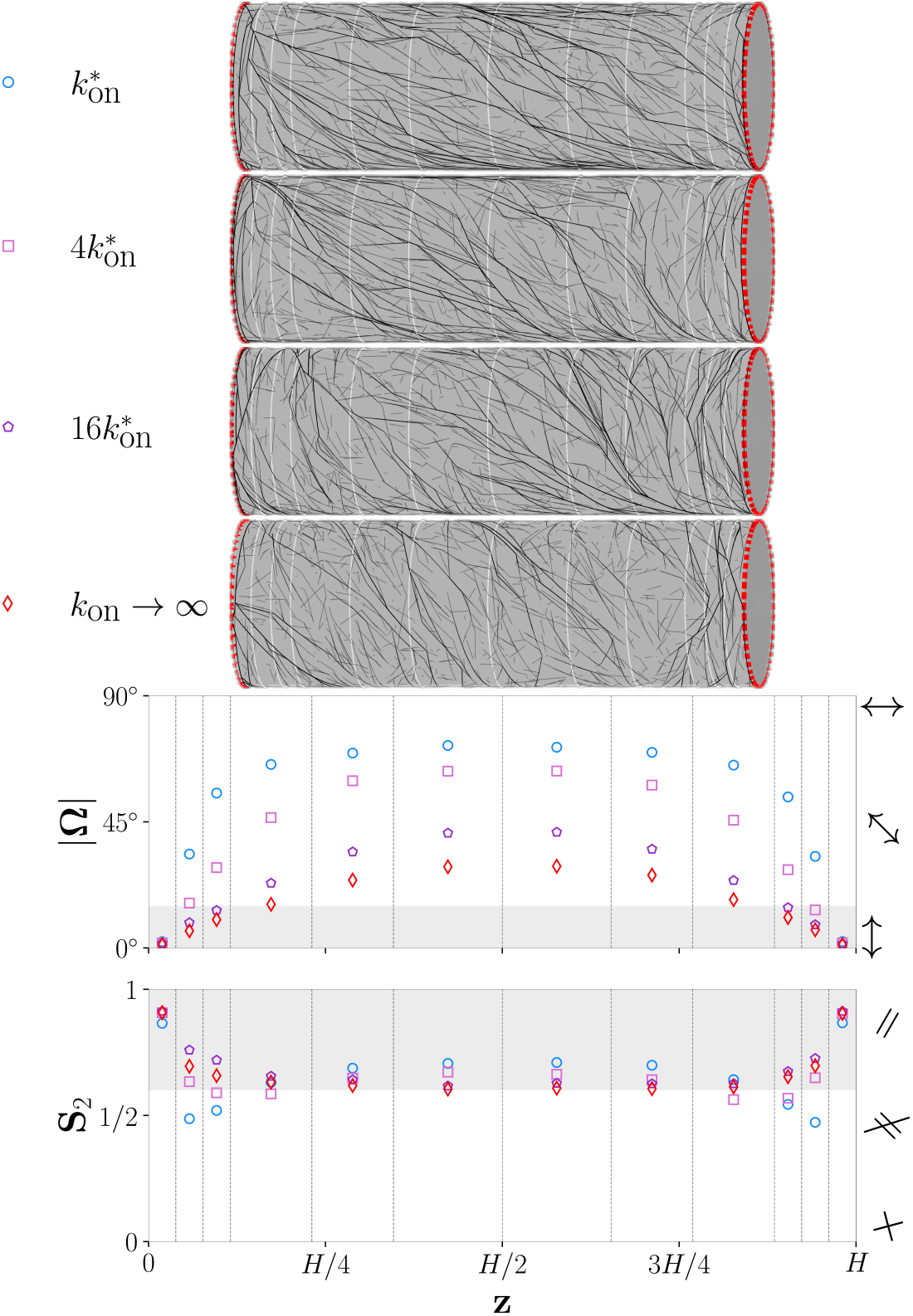
Ordering of cortical microtubule arrays in a long cell plotted for various values of the anchoring rate *k*_on_, with all other parameters set as default in Table S1. Top: visualization of individual simulations. Anchoring rates used were multiples of the experimentally estimated value 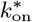, or taking the infinite (geodesic) limit. Bottom: averages of order parameters |Ω| (dominant angle) and *S*_2_ (order) plotted along the respective bands. Order parameter statistics are calculated by considering the angular distribution for microtubules within each band of the domain after 10h simulation time, and then averaging over 100 simulations. A detailed illustration of the distributions in the 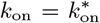 and *k*_on_ *→ ∞* cases is provided in Fig. 4, with the remaining cases (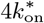 and 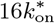) found in Fig. S11.

### Anchoring and EIC Cannot Reliably Recover Transverse Ordering in a Large Domain

Even after taking the limit to geodesics, thereby eliminating the deflection influence, EIC could not reliably recover a globally transverse array for the long cell geometry (see red diamond data points in the |Ω| plot of Fig 3). This showed that EIC had a weak effect on larger cells even in the most permissive case. We further investigated the local order parameters in the geodesic case by comparing the spread of the distributions with that of the MT deflection case in Fig. 4. Toward the middle of the domain for the geodesic case, we found *S*_2_ to be moderate with |Ω| having a large variance. This indicates that, farther from the edges, there is no strong preference for the orientation taken among different cells; organization arises randomly due to MT-MT interactions with a weak tendency toward the transverse direction. In the case of default anchoring, the longitudinal ordering was slightly more robust. This was seen by the smaller spread in order parameters away from the edges, and faster convergence in Figs. S8 & S9. The observation of weak transverse ordering due to EIC within larger domains, even in the geodesic limit, is consistent with previous observations of weaker order parameters [52, 30, 12, 25]. With the exception of a recent study done in parallel by Li et al. [30], previous studies to our knowledge have focused on determining the order parameters on a global scale, ignoring the possible effects of over/underestimating order parameters due to local variations such as EIC.

**Figure 4:**
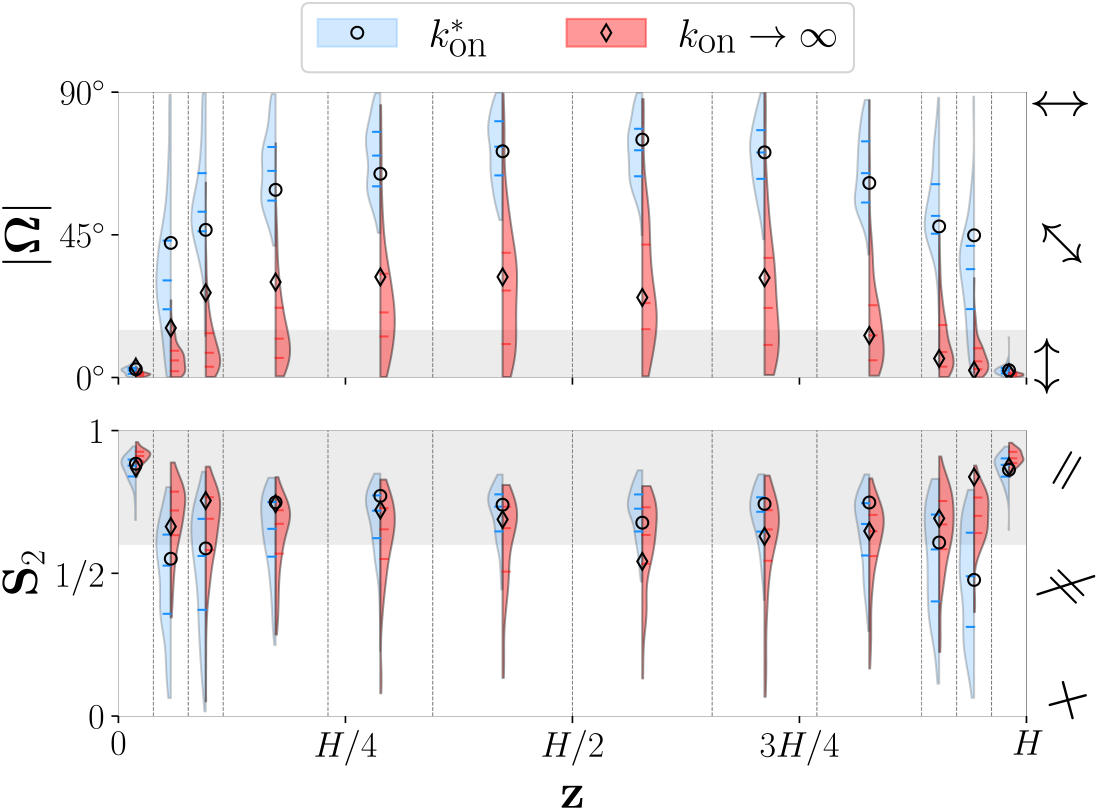
A closer examination of the ordering statistics in Fig. 3, using the same data. Violin plots are used to provide a qualitative visualization of order parameter distributions for the chosen anchoring rates. The circle and diamond indicators correspond to the order parameters of the respective simulation snapshots shown in Fig. 3.

It remains a possibility that the transverse arrays found in longer elongating cells are built on the existing array from when the cells were shorter. However, depolymerization experiments on already long elongating cells nonetheless recovered the transverse ordering from an initially empty or disordered state [23, 31, 60]. This is the same context in which our simulation takes place. We therefore examined a potential process to recover transverse arrays in the next section.

### In Long Cells, Orientation-Dependent Catastrophe Allows for Transverse Ordering Despite Longitudinal Deflecting Bias

We have shown that in the geodesic limit in long cells, global transverse ordering fails. With the addition of the longitudinal MT deflection, even short cells struggled to order transversely far from the catastrophe-inducing edges. The question remained as to how the transverse CMT ordering process can overcome the theoretical influence of MT deflection. A possible mechanism is the favouring of transverse MTs through an orientation-dependent regulation of dynamic instability – an idea that has been proposed as early as 1989 by Wasteneys and Williamson [60]. We tested this using a model in which MTs exhibit orientation-dependent catastrophe, for example, as a consequence of encountering the transverse texture created by cellulose microfibrils.

We introduced orientation-dependent catastrophe as described in the Methods section, where the parameter *ε* characterizes the intensity of angle dependence relative to intrinsic MT instability (see Eq. 3). In Fig. 5, we show that this global biasing mechanism overcame the longitudinal influence, with the default 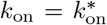, and created transverse arrays with a value of *ε* = 0.8. This corresponds to not quite doubling the catastrophe rate of longitudinal MTs relative to that of transverse MTs. This mechanism was robust in the sense that there was less spread in both |Ω|and *S*_2_ (Fig. S12) about the values of transverse ordering. It remains to be seen whether this is a realistic value that can arise from cellulose feedback and/or other potential sources of orientation-dependent catastrophe. It was also possible to control the sensitivity to the orientation-dependent catastrophe by increasing *k*_on_ (lowering the longitudinal influence). Seen in the bottom of Fig. 5, increasing *k*_on_ by a factor of four, thereby halving the mean first anchoring length to approximately 2.35 *µ*m, significantly lowered the *ε* values needed for transverse ordering. In this case, a 1.2 to 1.4 fold increase in catastrophe rate (*ε* = 0.2 *−* 0.4) was sufficient.

**Figure 5:**
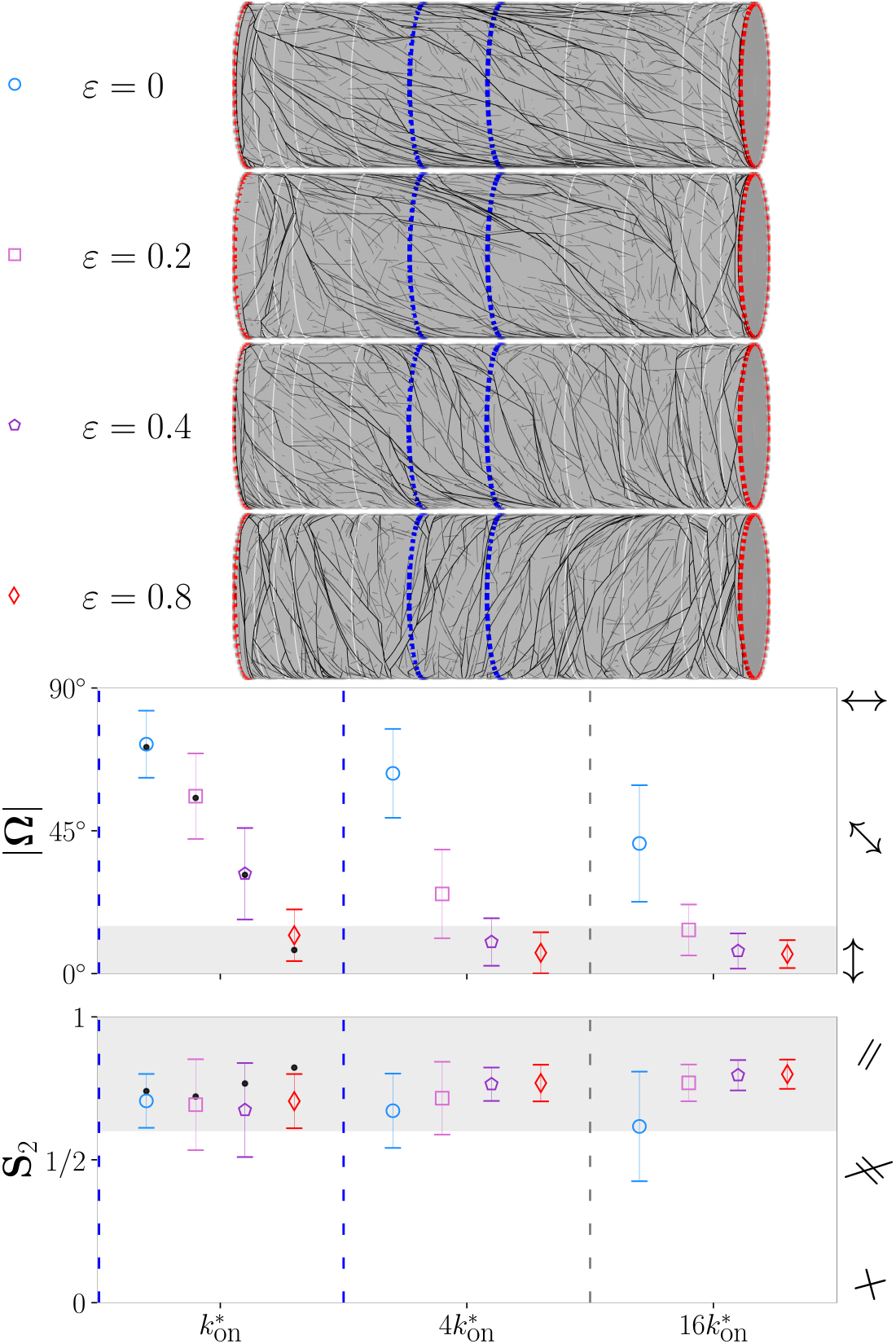
Ordering of the cortical microtubule array plotted for various values of orientation-dependent catastrophe strength *ε* (Eq. 3) and anchoring rate *k*_on_, with all other parameters set as default in Table S1 in a long cell. Top: visualization of individual simulations where *ε* is varied with fixed 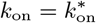. The middle-left band of each cylinder, highlighted with blue lines, indicates the region in which order statistics are calculated in the bottom plot. Bottom: order parameter statistics calculated in the middle-left band for chosen *k*_on_ values (represented by each column) and various *ε* values (represented by the same indicator shapes as above ordered from highest to lowest from the left in each column). Each shape and bar indicate the mean and standard deviation, respectively. Black dots in the *k*_on_ column (enclosed by the blue dotted lines) indicate the corresponding order parameters of each snapshot shown above. By symmetry, the ordering statistics are similar for the middle-right bands. Order parameters are calculated as in Fig. 3 with 100 runs except not all were 10h simulations since larger *ε* simulations organize quickly (see Supplementary Figs. S9 & S10), slowing the simulation down. Additional data for the 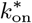 case are found in Fig. S12.

We note that, although the ordering parameters in the middle of the cell were similar for both 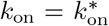 and 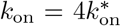 in the *ε* = 0 case (circle indicators in the first two columns from the left of Fig. 5 bottom subplot), the system 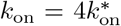 was more responsive to variations in *ε* (comparing e.g., the square indicators). This suggests that the longitudinal influence on order parameters saturates in the 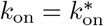 case.

We further observed that, for increasing *ε*, the convergence time of the mean *S*_2_ parameter was quicker than the geodesic case. With *ε* = 0.8, it was comparable to the speed of ordering due to edge-induced ordering in the bands closest to the edges (Fig. S10). This quicker convergence is consistent with the results in a recent study using a different computational method in the context of a three-state MT model, using *Arabidopsis* dynamic instability parameters[30]. There it was shown that, compared to other dynamic instability parameters, the modification of catastrophe rates has the most potent effect on both strength and speed of ordering in the particular settings of their simulation. This orientation-dependent catastrophe could be a factor needed to speed the ordering in simulations to times more accurate to those observed *in vivo*, in addition to the hypothesized role of MT-dependent nucleation [31, 42].

## Discussion

As CMT models are generalized to complicated geometries, one must consider whether previous assumptions remain applicable. In particular, the approximation of MTs travelling along geodesics has remained unchallenged. We present a model generalizing the previous MT-MT interaction modelling formalisms by explicitly modelling the mechanical deflection of MTs. This was done on a cylindrical cell, using biologically relevant parameters and dimensions. We have shown that such mechanics had significant impact, especially in long cells. The tendency of individual MTs to minimize bending energy translates into a strong influence on the collective array organization *in silico*. As a result, the array orients to the longitudinal direction. The presence of EIC can no longer be considered sufficient for the formation of transverse arrays throughout long cells. Modulating the degree of deflection via membrane-MT anchoring rate alone offers some control over this deviation from transverse arrays. In the case of short cells, EIC is sufficient to form transverse arrays, consistent with observations of the short cells in the root division zone of *clasp-1* mutants [3]. That transverse arrays can form in smaller cells provides an interesting hypothesis: is it possible that EIC produces the initial transverse array to kick-start the anisotropic expansion, after which there is a positive feedback loop from cellulose and/or stress/tension in the membrane? Observations in *Nitella* internodal cells indicate a correlation between the strain rate of elongating internodal cells and the prevalence of transverse MTs [38]. A model including an expanding domain may be of interest for future work.

Nonetheless, the observation of transverse CMT arrays recovering from an empty state after depolymerization in longer cells [60, 31] suggests the presence of additional processes to overcome the expected longitudinal bias. The necessity of additional processes is further illustrated by depolymerization experiments in *Nitella* [60]. In such a system, the domain is extremely large (5-15 mm in length for rapidly expanding cells, data from the experiments in [60]) compared to the width of the transverse boundary layer we observed. Although bending effects are expected to be weaker due to the larger radii of *Nitella* cells, one may expect EIC to be extremely weak simply due to the domain size. This is illustrated in Fig. 1 A, where there is an extreme difference in scale between the MT lengths and distance from the edge. Furthermore, CMT self-organization on the scale of *Nitella* cells solely due to MT-MT interactions may not be as robust. Yet, transverse CMTs manage to consistently recover. Potentially, an additional processes could involve mechanical forces shifting individual MTs toward the transverse direction. Such mechanisms could be incorporated by modifying the mechanical energy formulation for MTs. This could result in an effective deflection bias as implemented in [35], where the bias was given to MT tip orientation without an underlying mechanism. No observations thus far support the existence of such a process. Instead, we find this work to point toward an idea alluded to by Wasteneys and Williamson [60]: the existence of a “non-kinematic” process favouring transverse MTs through an orientation-dependent modification of dynamic instability. This is a reasonable hypothesis given the observed geometric effects of membrane shape and plethora of microtubule-associated proteins that are known to affect dynamic stability [3, 27, 61, 33, 37]. We have shown that an orientation-dependence of the catastrophe rate, such as through cellulose feedback, could result in robust transverse ordering despite the longitudinal bias induced by MT deflection. From our observation that the influence of orientation-dependent catastrophe gets stronger at higher anchoring rates, the sensitivity to such a mechanism could be controlled by the anchoring rate. The existence of such an orientational dependence may be tested by measuring the catastrophe rates of MTs at various angles with or without the presence of transversely oriented microfibrils. In addition, there is a long-standing hypothesis of MT mechanosensing [22, 21]. Currently, such a mechanism that is consistent with observations and physical principles is unknown, leaving any model largely phenomenological. We note that the orientation-dependent catastrophe hypothesis and the idea of MTs being sensors of mechanical stress or tension are not mutually exclusive. If membrane tension affects MT stability in a cylindrical cell, such a mechanism may introduce orientation-dependent catastrophe similar to what we introduced here. Membrane tension may influence other dynamic instability parameters (polymerization rate, rescue rate, etc.), giving rise to the more general idea of orientation-dependent dynamic instability. In earlier work, this orientation-dependent dynamic instability was proposed in a simplified continuum model of MTs [43]. More recently, such ideas were tested in a simple context of a flat simulation domain by Li et al. [30] by assuming a hypothetical linear relation between stress and MT dynamic instability parameters including, and not limited to, catastrophe. Their orientation-dependent expression is mathematically similar to our model. Although their method differed in model parameters (*Arabidopsis* vs. tobacco) and computational implementation (differing in algorithm and representation of MTs), it has nonetheless shown similar effects on CMT organization. This indicates a consistent need for such processes across plant models.

Our study focused on a minimal set up without severing or alternative modes of nucleation, which are known to impact MT organization *in silico* and *in vivo* [2, 17, 13, 12]. *In vivo*, severing has been implicated in assisting the process of CMT reorientation in response to blue light [32] and stress in the cell wall [54, 15]. *In silico*, severing has been shown to increase or decrease the ordering of CMT arrays depending on the location of severing and subsequent behavior of severed MTs. However, models of severing thus far have not shown a biasing effect on the overall CMT orientation (see e.g., [43]). Thus, we do not expect existing severing models to significantly impact the orientation of CMTs observed in this work.

Recent work on modelling a more realistic nucleation process [42] indicates that including such a nucleation process may decrease the sensitivity of CMT arrays to biasing cues and promote transverse arrays in the case of long cells with small radii. In the geometry studied presently, specifically cells with larger radii, we suspect this would not significantly affect the longitudinal influence. We hypothesize that such local density-dependent nucleation favours transverse arrays due to the smaller average-MT-length-to-circumference ratio. When this ratio is small, it is easier for transverse MTs to wrap around and approach their own minus ends – thereby creating denser arrays in this orientation. However, our work assumes a larger radius in comparison to average MT length. Indeed, from observations of CMTs in relatively large *Nitella* cells, it has been hypothesized that MT-dependent nucleation creates disorder in the CMT array [59]. Future work is needed to test the interplay between increased longitudinal bias (from MT deflection) and promotion of transverse arrays (from local density-dependent nucleation) at smaller radii.

Another degree of freedom unexplored in this work is the effect of changing the dynamic instability parameters aside from orientation-dependence, either as constants or time-varying. By decreasing the average MT length, which is governed by the activity of catastrophe factors [16], it might be possible to mitigate the deflection of individual MTs. More generally, hormones are known to affect MT organization via the regulation of microtubule-associated proteins; both their abundance and their localization in certain geometries. For example, brassinosteroids are known to regulate CLASP [39] and MDP40 [57] which affect MT stability, and the combination of auxin and gibberellic acid has been shown to reorient MT arrays where the proposed mechanism of action is the localization of MT nucleation complexes [55] to faces of *Arabidopsis* cells. Regardless, by keeping with a commonly used default parameter set [52, 13], our work serves to indicate the importance of this bending influence.

All CMT models thus far, including this model, have a common limitation: MTs are approximated as polymerizing along “fixed paths” where the MT body does not sweep laterally in response to mechanical forces. The fixed path approximation (described in Sec. S1.3) can be made to accurately model mechanical influences by considering the anchoring process; the accuracy relies on the average anchor length parameter *l*_*d*_ being small. Although this excludes the study of contexts where such movement is extreme, this model provides an approximation, a perturbation, of the fixed path model into a model of laterally dynamic MTs. The coupling of MT mechanics and anchoring provides a first step toward a more general model that has the potential to capture aspects such as the dynamic reorientation of CMT arrays. As applied to the context of bending mechanics, the observation of a “default” tendency for MTs to orient in the longitudinal direction [28, 5, 51] may be a factor in the transition from transverse to longitudinal arrays. This is a tempting hypothesis since this deflection is simply due to the physics of MT polymers; the cells would not need to expend additional resources for this to occur. Furthermore, it may be possible to incorporate the mechanics induced by cytoplasmic streaming, hypothesized to also play a role in reorientation [19, 40].

## S1 Supplementary Information

### S1.1 Microtubule Interactions

Upon a collision of an incident MT (Fig S1, black) and a barrier MT (Fig S1, grey), there are three possible outcomes: entrainment (incident MT follows the barrier MT), collision-induced catastrophe of the incident MT, or crossover (incident MT continues past the barrier MT) without severing. The outcome is determined by the angle of incidence: entrainment if below the critical angle Δ*φ*^*∗*^, and a 50% chance for catastrophe or crossover otherwise. This choice of critical angle is consistent with many of the models thus far [2, 3, 13, 17, 52, 35, 25, 45, 42], with some models differing in the probability of catastrophe [2, 3, 35, 25].

**Figure S1:**
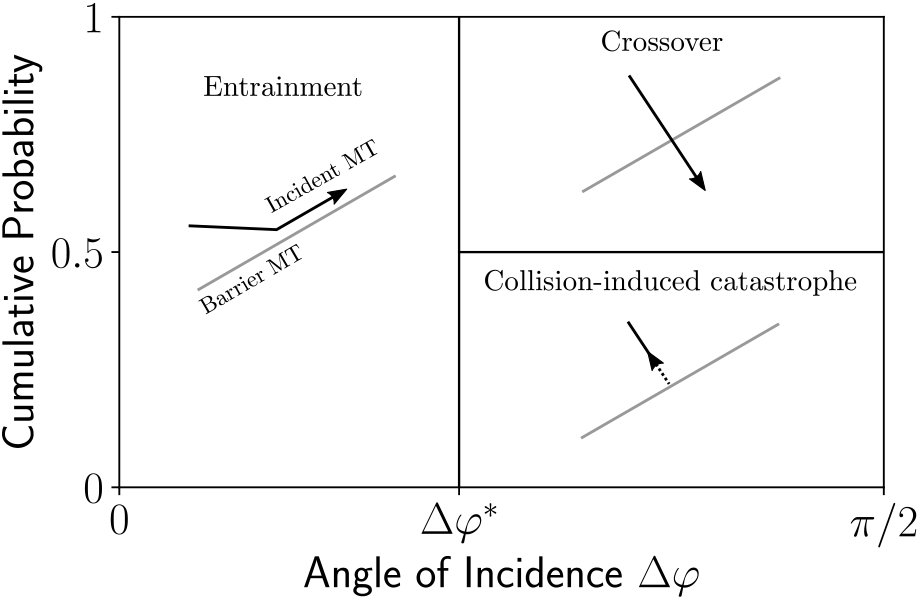
Illustration of collision resolution probabilities used. For angle of incidences less than 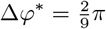between the barrier MT (grey) and the incident MT (black), the result is always entrainment. For larger angles, there is a 0.5 probability of collision-induced catastrophe or crossover. These values are taken from the model used in [2]. Illustration inspired and adapted from [52].

### S1.2 Table of Paramemters

**Table S1:** Parameters used in the simulation in dimensional form. Parameters are segregated in collections of rows from top to bottom describing: dynamic instability and nucleation rates, MT-MT interactions, cell geometry, and MT anchoring. We denote the default anchoring rate as 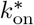, to distinguish from varying values of *k*_on_ used in the simulation. Catastrophe-inducing boundaries are at the top and bottom of the cylinder. Parameters from experimental measurements have been rounded, and others are estimated from casual observations. ^*∗*^ This catastrophe probability is a simplification of a more complicated probability distribution observed in [14].

| Parameter | Description | Value |
| --- | --- | --- |
| $v_g$ | Plus end growth speed [56] | $0.08 \mu\text{m s}^{-1}$ |
| $v_s$ | Plus end shrinking speed [56] | $0.16 \mu\text{m s}^{-1}$ |
| $v_t$ | Minus end shrinking speed [46] | $0.01 \mu\text{m s}^{-1}$ |
| $r_n$ | Nucleation rate [17] | $0.001 \text{ s}^{-1} \mu\text{m}^{-2}$ |
| $r_{\text{obs}}$ | Observed catastrophe rate (see Methods) [56] | $0.0045 \text{ s}^{-1}$ |
| $r_r$ | Rescue rate [56] | $0.007 \text{ s}^{-1}$ |
| $\Delta\varphi^*$ | Critical angle [14] | $40^\circ$ |
| $p_{\text{cat}}$ | Catastrophe probability [14]* | 0.5 |
| $p_{\text{zip}}$ | Zippering probability [14] | 1 |
| $R$ | Cylinder radius | $20 \mu\text{m}$ |
| $L$ | Cylinder length | $2\pi R$ (long) or $2R$ (short) |
| $p_{\text{edge-cat}}$ | Probability of edge-induced catastrophe | 1 |
| $k_{\text{on}}^*$ | Default anchor rate [1] | $0.0057 \text{ s}^{-1} \mu\text{m}^{-1}$ |
| $l_d$ | Default mean first-anchoring length (dimensional) | 0.24R |

### S1.3 Simulation Method: Individual MTs

We independently implement the event-driven simulation method outlined in [52] and generalize to the case of curved MTs. As the length of a MT changes, it “sweeps” through various mechanical equilibria (energy minimizers), driving a lateral movement. For illustration purposes, we refer to the top of Fig. 1 B where this sweeping is visible as the length increases. All models thus far approximate MTs as following paths without sweeping; we term this the “fixed path approximation”. A model with sweeping MTs is computational challenging; we therefore also make use of a fixed path approximation. Coupling this with the anchoring process allows us to capture MT deflection with control on the various errors introduced. We provide a brief explanation below; a thorough analysis of this approximation is the subject of separate manuscript in preparation.

For this section, we assume a non-dimensionalization where length is in units of the cylinder radius *R* and time is in units of time required for polymerization to grow a MT to a length of *R*. The deflection of MTs only depends on MT lengths relative to the radius, and the speed is such that a MT will grow to a length *t* after *t* time units. Effectively, we have set *R* = 1 and *v*_*g*_ = 1. We parametrize the MT path with its tangential angle as a function of arc-length and time, *φ*(*s*; *t*), resulting from the variational problem. Our fixed path approximation comes from a spatio-temporal approximation on the MT shapes. We describe it in Sec S1.3.1 and S1.3.2, with an accompanying illustrative summary in Fig. S2.

#### S1.3.1 Step 1: Temporal Approximation

Suppose we seek to simulate a MT of length *L*. We first approximate the MTs as following curves that are fixed in time. Although MT segments eventually become fixed in shape due to the anchoring model, their shapes nonetheless sweep as they grow prior to being fixed (as described by the solutions to Eq. 1). Referring to the visual in Fig. 1 B, the growing MT sweeps through the various green curves before finally reaching the the longest green curve. We approximate the MT growing along the longest green curve rather than simulating each smaller curve as time progresses. Mathematically, the approximation made is,

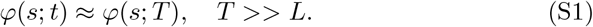

Here, *T* is a time after all anchors have fixed the MT shape up to length *L*. This is illustrated in Step 1 of Fig. S2. In this case, we are only interested in the MT for times before the first anchor, so *T > t*_4_ is the time after the first anchor has fixed the shape. Although visibly inaccurate in the exaggerated case of Fig. S2, this approximation can be justified in the case where *l*_*d*_ is sufficiently small. In such cases, MTs do not sweep significantly on each individual segment. This approximation only applies to the freely growing tip. Furthermore, in the case of small *l*_*d*_, the segments of barrier MTs will likely be already anchored prior to collisions, meaning the inaccuracies in collision geometry caused by this approximation are reduced.

#### S1.3.2 Step 2: Spatial Approximation

We now construct a numerical approximation to the curve *φ*(*s*; *T*). First, we solve the variational problem for *φ* numerically using a boundary value problem solver with a fine grid size *δs*. It remains to coarsen the spatial discretization of the curve for the purposes of resolving collisions in a computationally efficient manner. This is because MTs will be approximated as line segments, and many segments result in increased calculations for collision detection. We set a coarse discretization 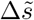 that provides an upper bound on the segment lengths used in the simulation. Suppose there is an anchored segment with anchor points *s*_1_ *< s*_2_. Then, the discretization represents the actual path 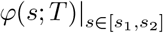using piecewise linear segments of lengths Δ*s*_*i*_ with corresponding subdivision points 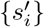 on the interval [*s*_1_, *s*_2_]. If there are *n* segments, these segment lengths are given by,

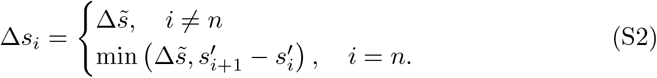

In words, all segments have 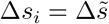 except for the last one where it is possibly a remainder that is less than 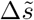. We choose to write the coarse segment angles 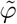 as the average,

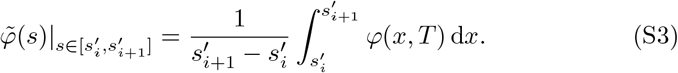

This coarsening method is chosen such that the length of the MT is preserved, different from e.g., an interpolation method. A result of this is that the coarsened linear approximation is not necessarily composed of secants or tangents on the original curve, as illustrated in Step 2 of Fig. S2. There is no longer a dependence on time *t* since the MT now has a fixed path up to the length of interest. With these values for MT angles, we represent MTs by mapping to cylindrical coordinates.

**Figure S2:**
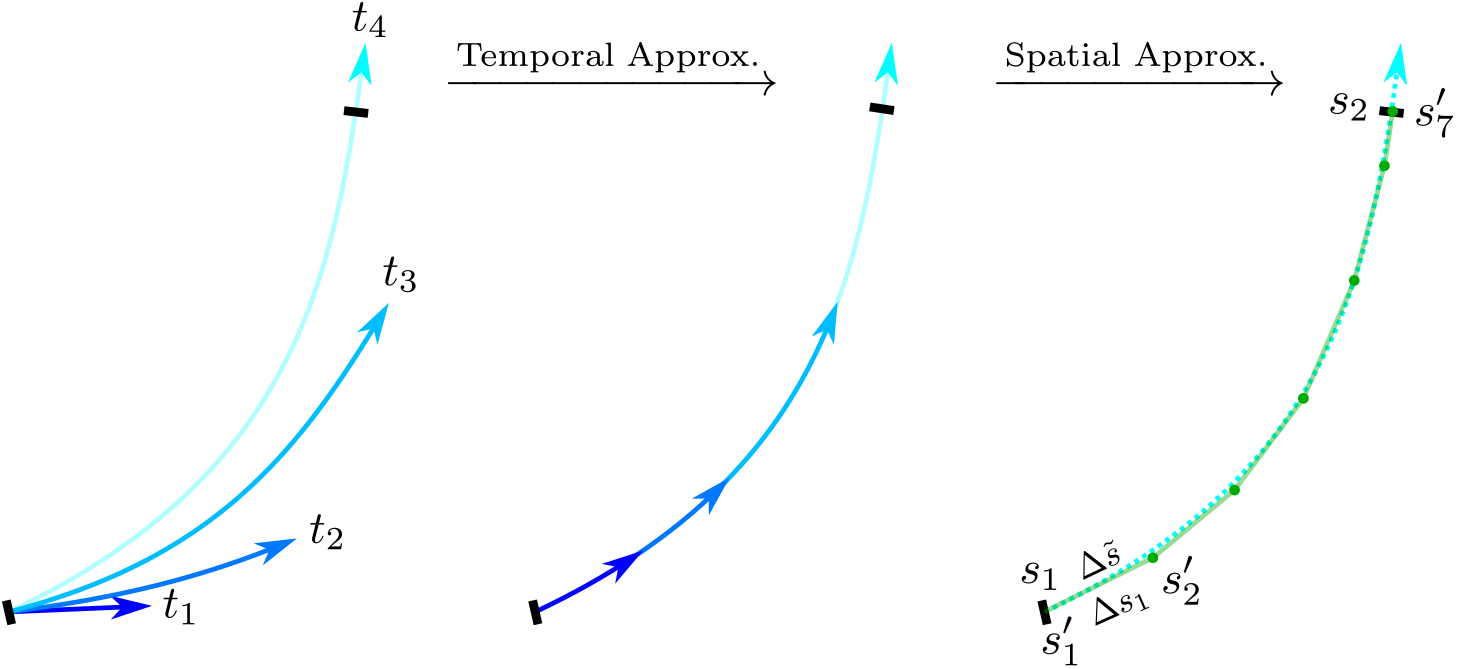
Approximating MT paths as fixed paths. Temporal approximation: the MT body (shades of blue on the leftmost curves) sweeps as it grows. For times before the anchor time *t*_4_, the MT is approximated as tracing the future anchored segment (light blue corresponding to *t*_4_) in Eq. S1. Spatial approximation: the path is spatially discretized (green polygonal curve) as in Eq. S3. Labelled are the anchor points *s*_*i*_, discretized subdivision points 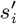, upper-bound segment length 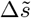, and implemented segment lengths Δ*s*_*i*_. Note the spatial approximation is not simply an interpolation i.e. the segments (translucent green) are not secants of the previous curve (dotted blue), see Sec S1.3.2.

In the present work, the fine grid solution is solved with *δs* = 10^*−*2^ using SciPy’s `ode.solve_bvp` with a tolerance of 10^*−*4^. The tolerance controls the accuracy of the solution, whereas the fine grid determines the desired points of the solution to solve for. Then, the coarse spacing is set as a function of initial MT angle *φ*_0_. Without loss of generality, we restrict to *φ*_0_ *∈* [0^*°*^, 90^*°*^],

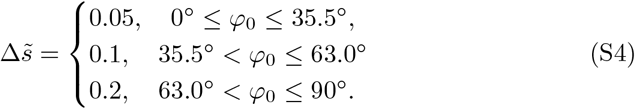

The reasoning for the piecewise values is because solutions with initial angles close to 90^*°*^ do not vary much. Therefore, we do not need 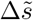 to be small. The choice of where to divide the piecewise function is otherwise arbitrary.

We find this choice to give an average approximation error to be less than 1^*°*^ on individual MT shapes. We leave further error analysis to a manuscript in preparation.

### S1.4 Simulation Method: Interacting MTs

Conceptually, the simulation algorithm follows that described in [52]. There are deterministic and stochastic events. The deterministic events are collisions between the line segments representing MTs whose times are calculated given the current state of the system. The stochastic events are sampled using a kinetic Monte Carlo (Gillespie) algorithm. All events are sorted by their times, and the earliest is executed. Upon execution, the appropriate changes in simulation state are made and new events are calculated and sampled. This is repeated.

In addition to the usual interactions described in Sec. S1.1, the introduction of MT bending requires new decisions on MT dynamics within bundles. As with [52], bundles are computationally represented as occupying the same curve such that collisions with bundles are equivalent to collision with a single MT. We identify bundles as a partition on the set of MTs, making “bundling” an equivalence relation on this set. Being able to distinguish between bundles allows us to resolve collisions between entrained MT bundles, as illustrated in Fig. S3. We keep track of the “sidedness” of entrained MTs, such that we can resolve the collision illustrated in Fig. S3b). In this sense, our simulation captures elements of the simulation methods in [2, 3, 35] where the non-zero spacing of entrained MTs is explicitly represented. This differs from the choice made in [52] and other studies based on the methods described therein [53, 13]. Our method also allows for ignoring MT “sidedness”, and we note that this generally gives similar results albeit with higher ordering *S*_2_. The question of which choice is most appropriate remains open; this relies on more precise measurements on the nature in which MTs bundle.

**Figure S3:**
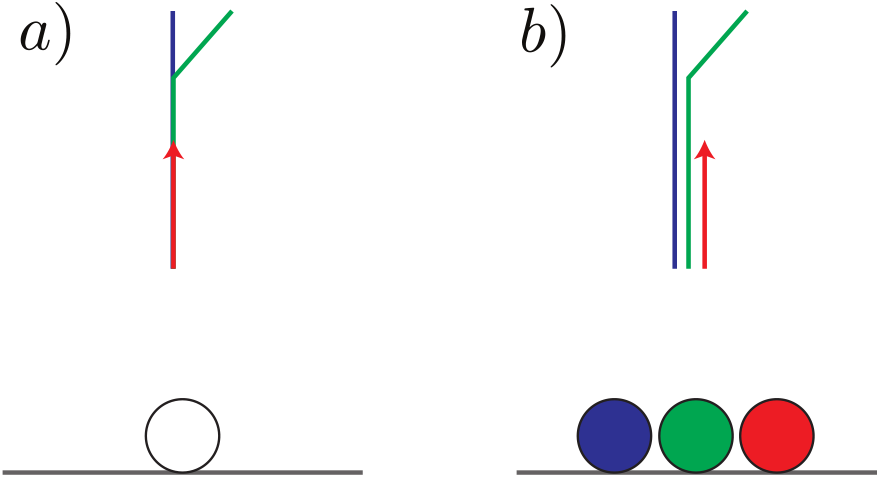
Representation of bundles within the simulation. a) Entrained MTs are represented as occupying the same curve. b) illustrates an alternative way of representing bundles where entrained MTs do not occupy the same curve. In the case of b), the leftmost red MT is able to resolve the collision with the entrained centre green MT. Although our simulation uses representation a), we keep track of the “sidedness” of entrained MTs, maintaining one of the benefits of b).

Lastly, there are new events associated with the “freeing” of bundled MTs, whereby an entrained MT is no longer entrained. Once this occurs, the liberated MT is allowed to deflect, as illustrated in Fig. S4.

**Figure S4:**
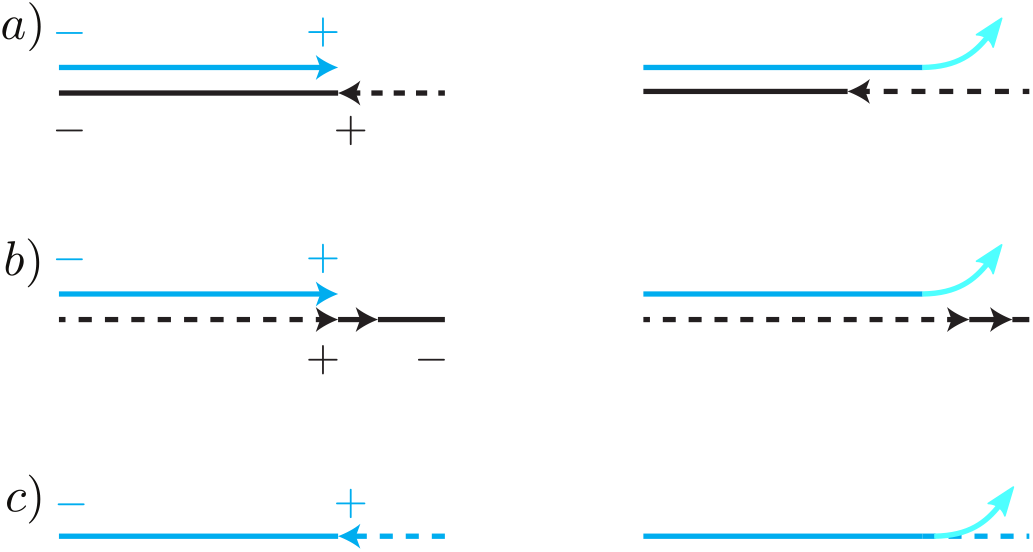
Illustration of how a MT of interest (blue) may branch out of its entrainment (black) through: a) blue MT catching up to the shrinking black MT, b) the black MT shrinking faster than the blue MT is growing, c) the blue MT is shrinking but rescued. In all these cases, the blue MT tip is freed and there is a new path (light blue) because there is a new sequence of anchors starting from its growing tip and hence a potentially different deflection. This is as opposed to the blue MT re-tracing the path it was previously was on.

### S1.5 Order Parameters computed in bands

The order parameter *S*_2_ from [13] is used,

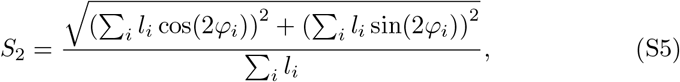

where *l*_*i*_ are the segment lengths of each MT with corresponding angles *φ*_*i*_. The angle of *φ*_*i*_ = 0^*°*^ or *φ*_*i*_ = 180^*°*^ corresponding to the circumferential direction. This is derived as the positive eigenvalue of a tensorial order parameter. It measures order where each angle is weighted by the length of their respective segment lengths, and is equivalent to that used in [2]. The order values range from 0 (completely isotropic) to 1 (completely aligned), and is invariant of MT polarity. The dominant direction, Ω, is the orientation of the corresponding eigenvector, given as,

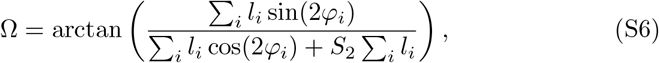

for segment lengths and angles *l*_*i*_, *φ*_*i*_ as in Eq. S5. The range of Ω is given by the interval (*−*90^*°*^, 90^*°*^), reflecting the invariance under polarity.

In order to study how behaviour varies near and far from the end caps, these order parameters are calculated for MT populations along bands of the cylinder, and each band’s order parameters are averaged among the respective order parameters over *N* runs. We use the usual average

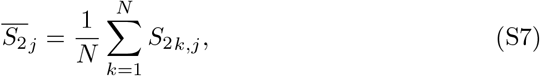

where 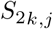 denotes the order parameter in the *k*th simulation, in the *j*th band. This index notation applies similarly to Ω_*k,j*_. However, Ω_*k,j*_ is averaged differently; we first take the absolute value, and then average:

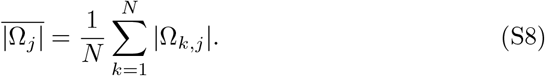

This is because directly averaging could result in 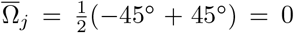, which would be misleading because both angles are equally diagonal.

### S1.6 Normalization of Catastrophe Rates

For this section, the angles are measured in radians. The previously measured catastrophe rate does not take into account the possibility of angle dependence, and we seek to normalize *r*_*c*_ with respect to *r*_obs_. Given *N* (*t*) growing MTs experimentally sampled at a particular time, the total observed catastrophe rate is *N* (*t*)*r*_obs_. If the angular distribution of the sampled MTs is represented by the number density, 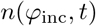 where 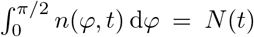, we set the equality,

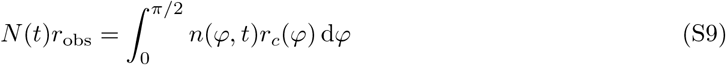

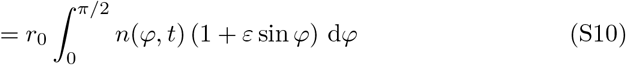

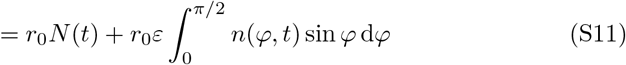

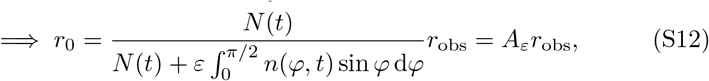

where we have defined *A*_*ε*_ to be the factor modifying *r*_obs_. Equivalently, one can view the above equality as a restatement of calculating the catastrophe rate as the average over the samples,

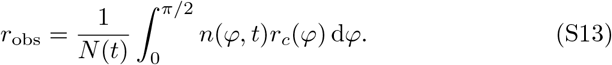

With this, we normalize to the instance in time in which *r*_obs_ is measured. Under the assumption that at the time of sampling, the sampled distribution was uniform, 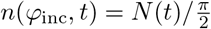, we can solve Eq. S12 to get,

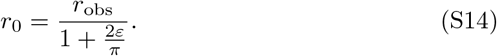

The catastrophe rate used here was measured in interphase cells [56] where the assumption of transverse cellulose microfibrils likely holds. Since these previous experiments did not take into account the orientation of MTs when measuring *r*_obs_, there is little data to indicate the orientations, and therefore the angular distribution (*n*(*φ*_inc_, *t*)), of the observed MTs. In the absence of further information, the assumption that MTs were chosen randomly without preference to a particular orientation is a guess.

Other possibilities of *n*(*φ*_inc_, *t*) are if only oblique MTs (defined to be *φ*_inc_ = *π/*4 for convenience) were sampled, or less likely, only longitudinal MTs. Transverse MTs are more difficult to observe since they are parallel to the array. To examine how *A*_*ε*_ (Eq. S12) changes depending on the sampled distribution *n*(*φ*_inc_, *t*), we plot *A*_*ε*_ as a function of *ε* under the assumption of a uniformly sampled MT distribution, and Dirac delta functions where only MTs of a particular angle are sampled. The angles chosen for the Dirac delta functions correspond to only longitudinal (*φ*_inc_ = *π/*2) or only oblique MTs (*φ*_inc_ = *π/*4). This is shown in Fig.S5. For the *ε* values used in our study, the differences are not large (especially for measuring oblique MTs).

**Figure S5:**
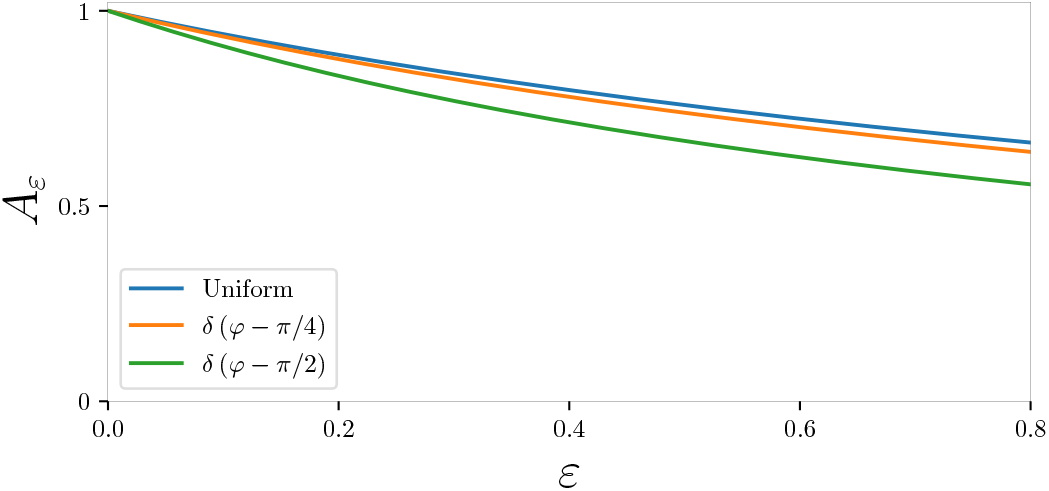
The normalization factor *A*_*ε*_ in Eq.S12 as a function of angle-depence strength *ε*, with different assumptions on the sampled MT number density *n*(*φ, t*) (coloured lines).

### S1.7 Examining Geometry and Microtubule Parameters

We investigate dependence of transverse boundary layer ordering and global longitudinal ordering on the domain size, anchoring rate, average MT length, and collision-induced catastrophe probability. The results are shown in Fig. S6 & S7.

#### S1.7.1 Boundary Layer and Anchoring Rate

We first quantify the boundary layer depth as the anchoring rate is varied in Fig. 3. We fit (`scipy.optimize.curve_fit`) the mean dominant angle |Ω| on half the domain to a Hill function of the form,

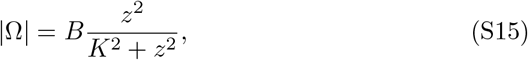

where *z* is the height as measured from the left-end, *B* describes the maximum value, and *K* describes the location of the half-max. The location of the half-max indicates how far the boundary layer propagates within the domain. While it is visually clear that the maximum value decreases as the anchoring rate increases in the top figure of Fig. S6, it is less clear whether the half-max changes its location. Plotting the fitted *K* values, we find that the half-max travels further within the domain as the anchoring rate is increased. This confirms that the effect of EIC propagates further (on average) as anchoring rate is increased.

**Figure S6:**
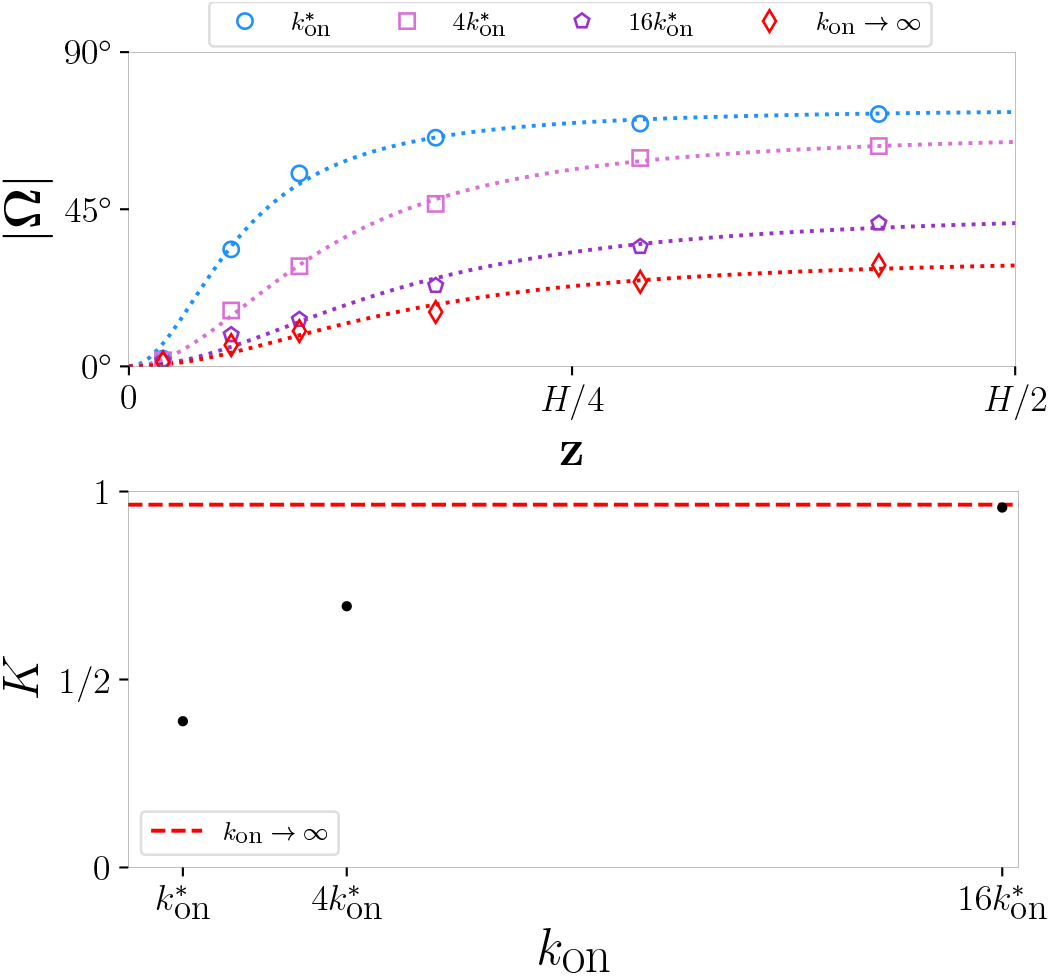
Quantifying the boundary layer shapes observed in Fig. 3 as the anchoring rate *k*_on_ is varied. Top: the mean dominant angles are fitted to a second-order Hill function of the form in Eq. S15. Bottom: the values of the half-max *K* (non-dimensionalized by the radius) from the fitted functions above, for each anchoring value. Dotted red line corresponds to the value from the geodesic limit.

#### S1.7.2 Boundary Layer and Cell Size

From Fig. S7, we find that, when MT bending is present with 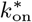, beyond a certain cell height (between 50 *−* 60*µ*m), there is a steep transition from localized transverse MTs to longitudinal MTs. The shape of this transition, and consequently the size of the boundary layer, appears to asymptotically become constant as the domain increases.

#### S1.7.3 Boundary Layer and Average MT Length

Since there is no analytic expression for the average MT length in this system of interacting MTs, we denote the “average length” as that for non-interacting MTs, given by [52]:

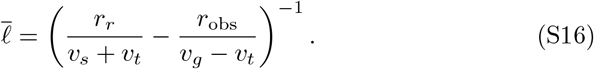

By simultaneously multiplying *r*_*r*_ and *r*_obs_ by the same factor, we change the average length by the same inverse factor. Therefore, in the lower-left pair of subplots Fig. S7, we find increasing the average MT length by 3*/*2 to have little discernible effect on the ordering observed previously. In the lower middle plots, we find decreasing the average MT length by 3*/*4 to slightly increase the dominant angle within the boundary later.

#### S1.7.4 Organization with Weak Interactions

By decreasing the probability of collision-induced catastrophes in steep collisions to *p*_cat_ = 0.1, we find ordering to be generally weak except at the edge of the domain. The dominant angle is biased toward the longitudinal direction but with large variance. This suggests that the longitudinal influence on bending individual MTs is not strong enough to produce a sharply peaked orientation distribution and that a stronger longitudinal influence arises through the collective behaviour of interacting MTs. On the other hand, the transverse influence of the edges is also significantly weakened with weaker interactions.

**Figure S7:**
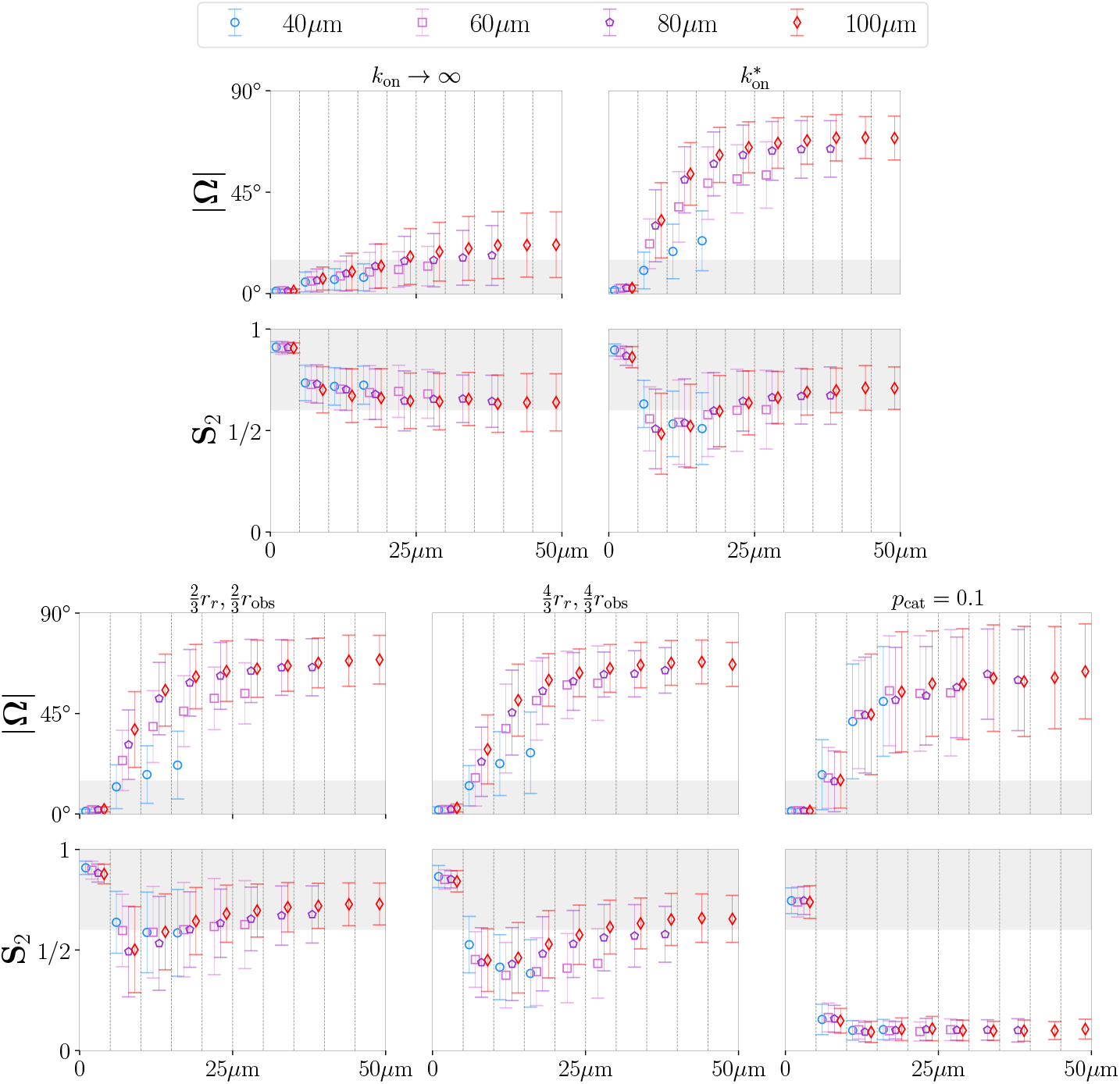
MT array ordering with differing geometries and parameters. Local order parameters are plotted in a similar manner as Fig. 3, taken over 100 runs after 10h simulation time. Each vertical pair of *S*_2_ and |Ω| plots represents results for a single parameter set where only the domain length is varied. The title above each pair of plots indicates the parameter(s) that have been changed from the default set, other than the domain length. Within each parameter set, the length of the cell is varied up to the maximum 100*µ*m, represented by the indicators plotting the mean and standard deviation. For each parameter set, the order parameters are only plotted for half of each domain size. For this reason, the there are fewer data points for the shorter domains. Due to symmetry, the other half is statistically similar.

### S1.8 Additional Plots

**Figure S8:**
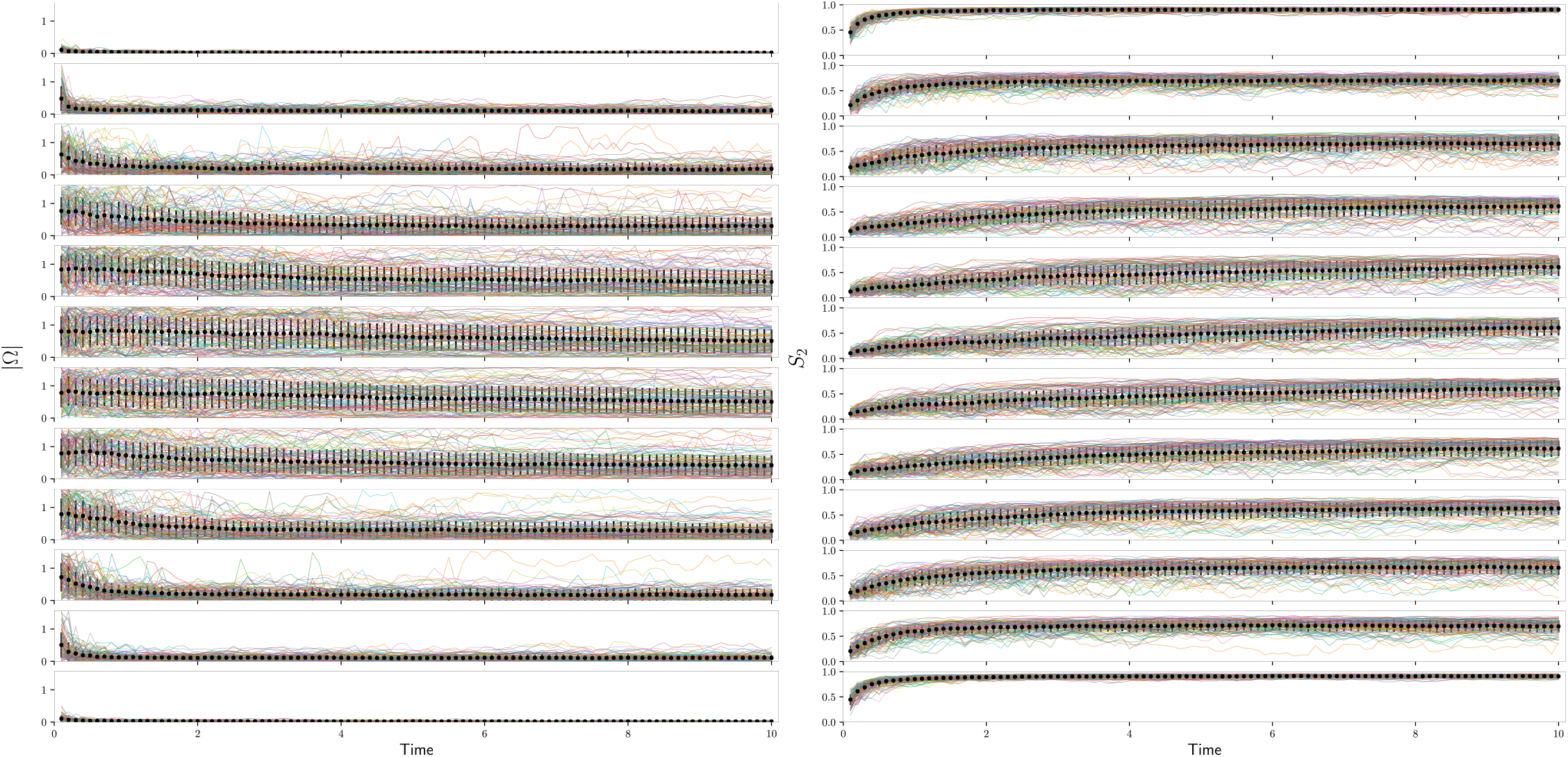
Order parameters over time for long cells, geodesic case and without orientation-dependent catastrophe. Each plot is split into the respective bands of the cylinder, partitioned in the same way as Fig. 3. The top to bottom ordering of each band here corresponds to the left to right ordering of each band in Fig. 3. Angles are measured in radians. 100 simulations are shown.

**Figure S9:**
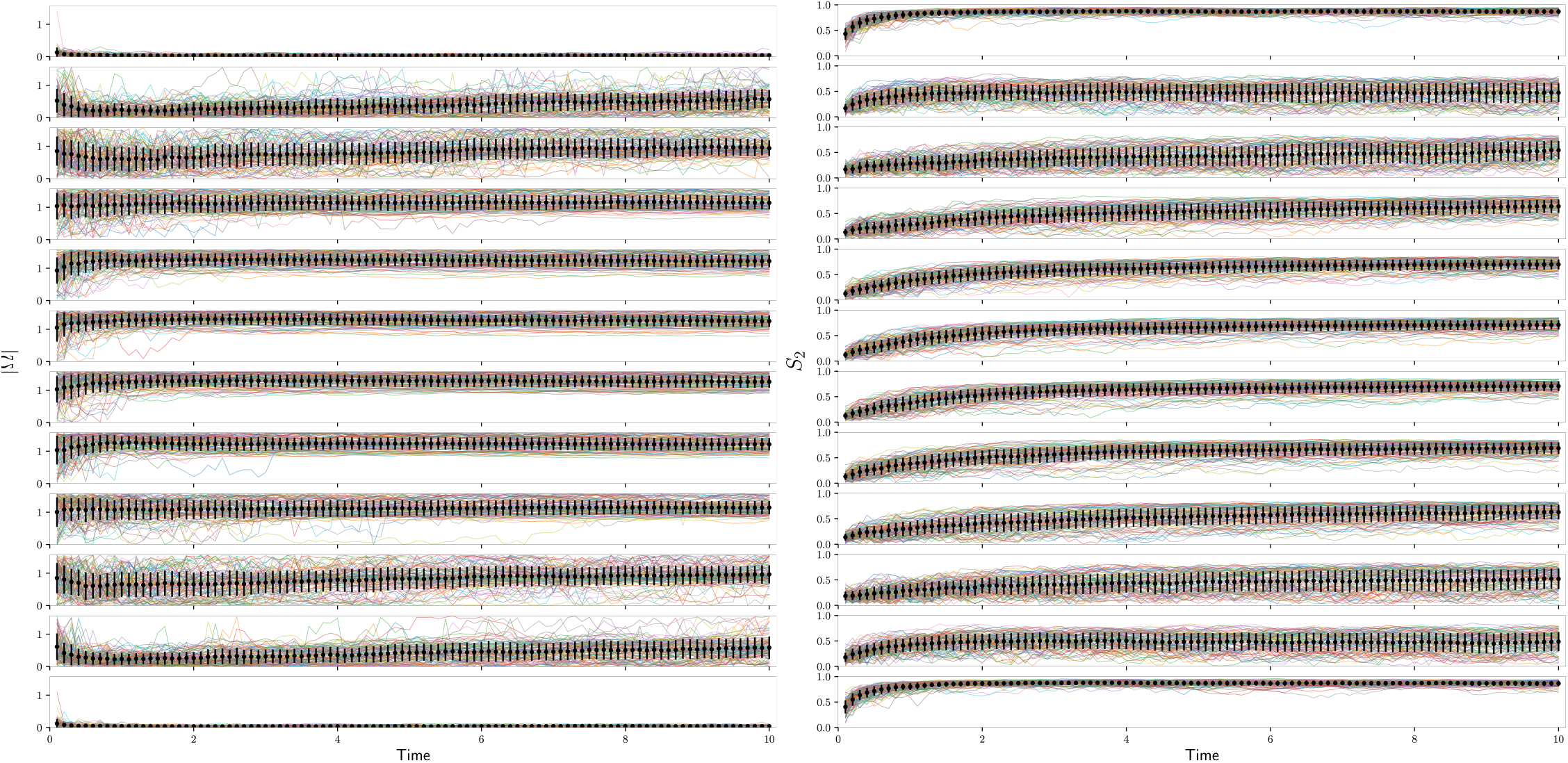
Order parameters over time for a long cell with 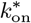 and without orientation-dependent catastrophe. The cylinder is partitioned in the same way as Fig. 3. 100 simulations are shown.

**Figure S10:**
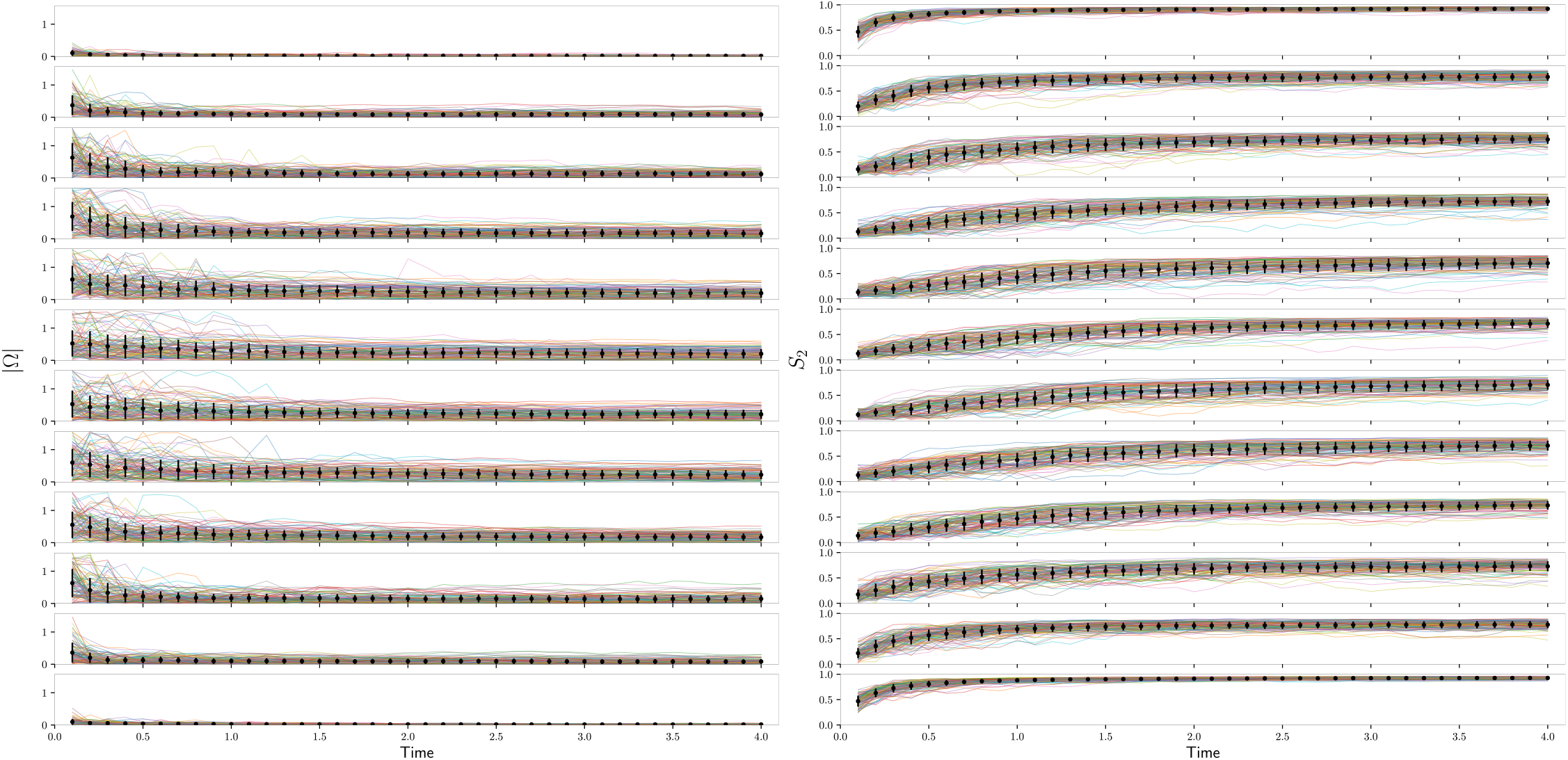
Order parameters over time for a long cell with ε = 0.8 and 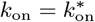. The cylinder is partitioned in the same way as Fig. 5. 100 simulations are shown.

**Figure S11:**
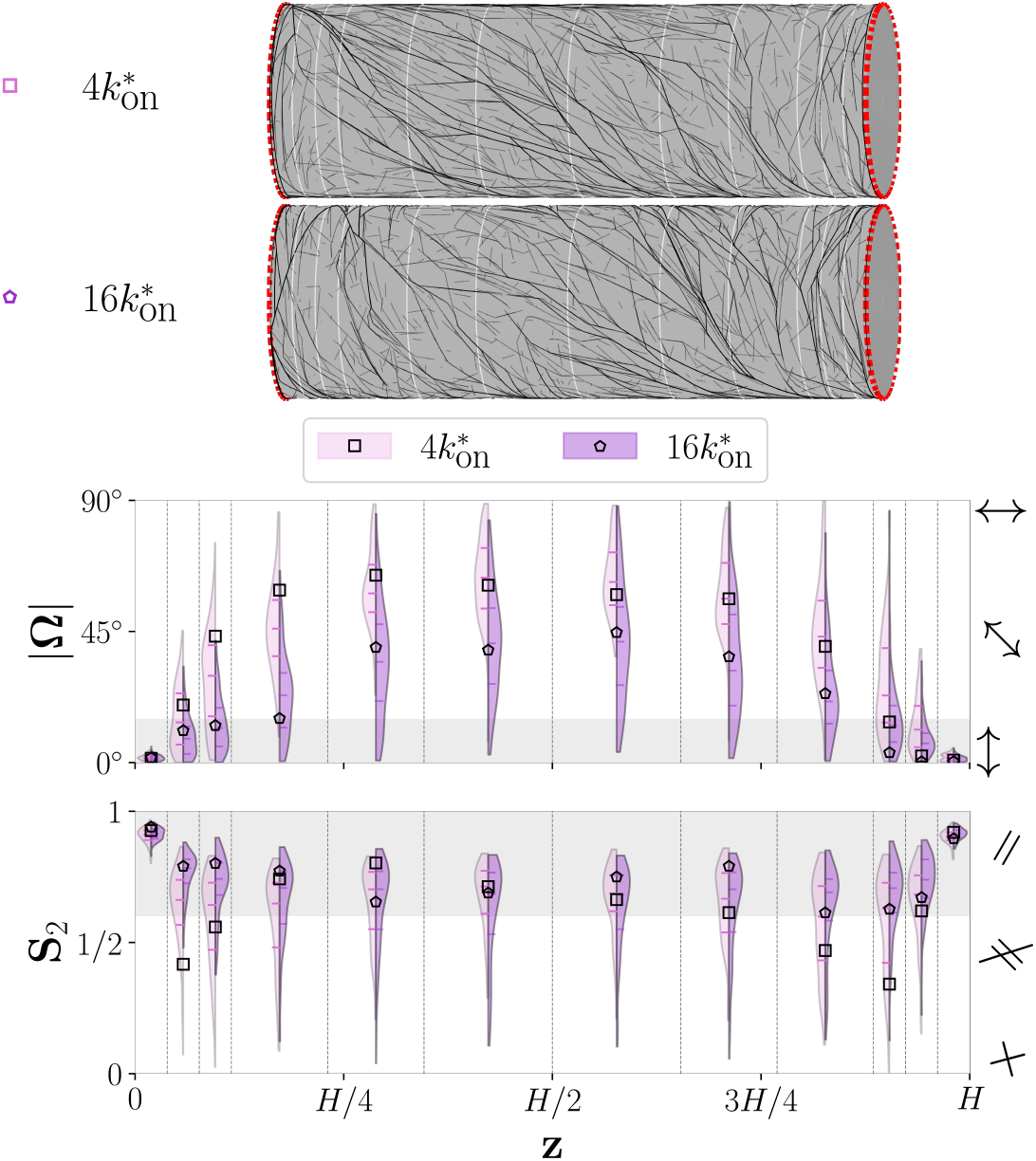
A closer examination of the ordering statistics in Fig. 3, using the same data over 100 simulations. Top: the same images of simulated MTs with the corresponding anchoring rates in Fig. 3. Bottom: violin plots for the distributions of order parameters, with the square and pentagon indicators representing the order parameters for the specific simulations snapshot shown at the top.

**Figure S12:**
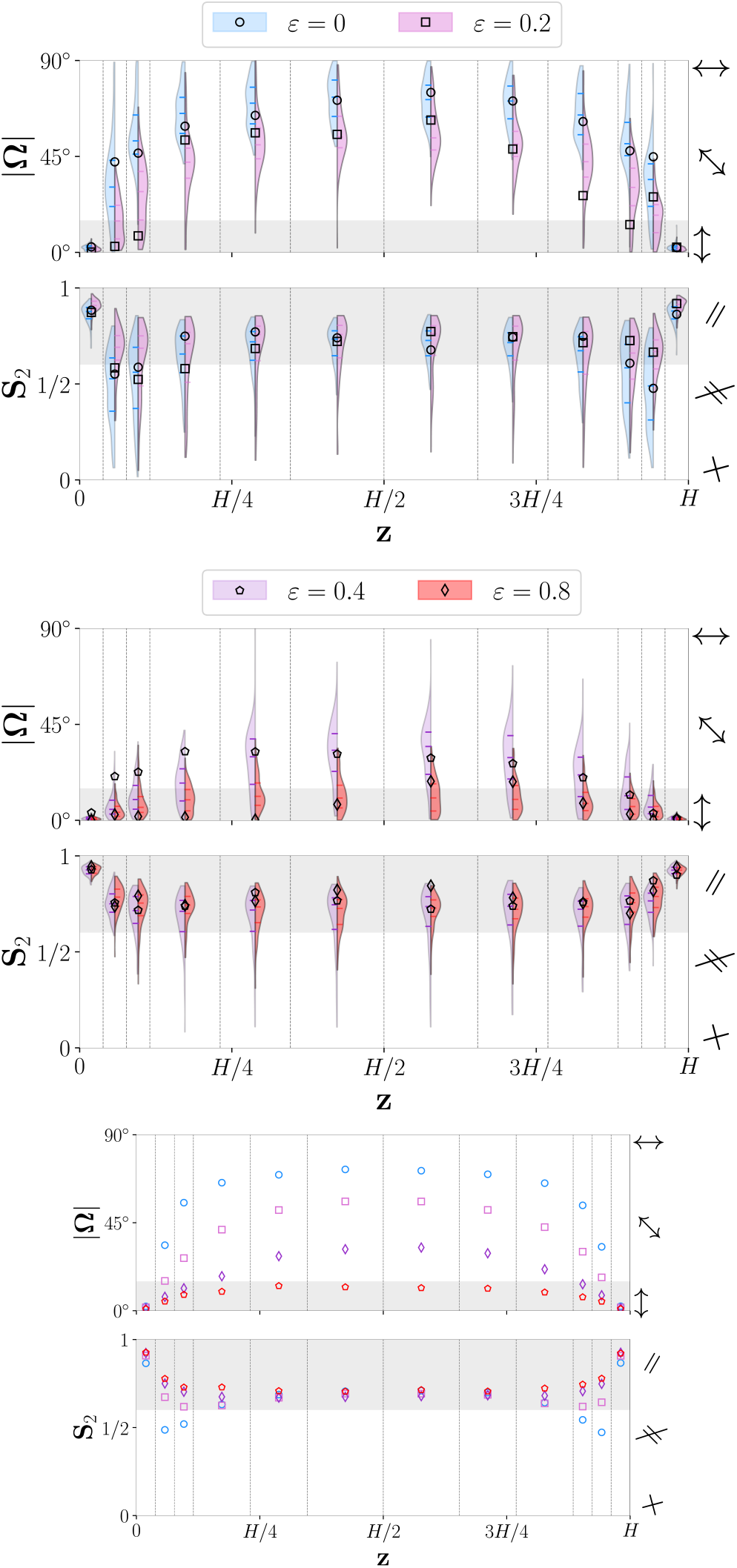
A closer examination of the ordering statistics in Fig. 5, using the same data over 100 simulations. The cylinder is partitioned in the same way as Fig. 5. Top four plots: violin plots for the distributions of order parameters, with the shape indicators representing the order parameters for the specific simulations snapshot shown at the top of Fig. 5. Bottom two plots: the mean order parameters for each *ε* value. The coloured indicators here represent the mean values of the distributions shown above.

## Notes

### Competing Interest Statement

The authors have declared no competing interest.

### Summary of Updates

Revisions in grammar and wording throughout; new analysis added in results and supplementary.

### Funding

- NSERC

